## Supplementary material for "Spatio-temporal mechanisms of consolidation, recall and reconsolidation in reward-related memory trace": Manuscript LaTeX source: catfish.bib

@ARTICLE{Wang2006,
author={Wang, K.H. and Majewska, A. and Schummers, J. and Farley, B. and Hu, C. and Sur, M. and Tonegawa, S.},
title={In Vivo Two-Photon Imaging Reveals a Role of Arc in Enhancing Orientation Specificity in Visual Cortex},
journal={Cell},
year={2006},
volume={126},
number={2},
pages={389-402},
doi={10.1016/j.cell.2006.06.038},
}

@book{Muller1900,
  title={Experimentelle beitr{\"a}ge zur lehre vom ged{\"a}chtniss},
  author={M{\"u}ller, Georg Elias and Pilzecker, Alfons},
  volume={1},
  year={1900},
  publisher={JA Barth}
}
@article{Lechner1999,
  title={100 years of consolidation—remembering M{\"u}ller and Pilzecker},
  author={Lechner, Hilde A and Squire, Larry R and Byrne, John H},
  journal={Learning \& Memory},
  volume={6},
  number={2},
  pages={77--87},
  year={1999},
  publisher={Cold Spring Harbor Lab}
}
@article{Alberini2006,
  title={Memory: mechanisms of memory stabilization and de-stabilization},
  author={Alberini, CM and Milekic, MH and Tronel, S},
  journal={Cellular and Molecular Life Sciences CMLS},
  volume={63},
  pages={999--1008},
  year={2006},
  publisher={Springer}
}
@article{Squire2015,
  title={Memory consolidation},
  author={Squire, Larry R and Genzel, Lisa and Wixted, John T and Morris, Richard G},
  journal={Cold Spring Harbor perspectives in biology},
  volume={7},
  number={8},
  pages={a021766},
  year={2015},
  publisher={Cold Spring Harbor Lab}
}
@article{Bontempi1999,
  title={Time-dependent reorganization of brain circuitry underlying long-term memory storage},
  author={Bontempi, Bruno and Laurent-Demir, Catherine and Destrade, Claude and Jaffard, Robert},
  journal={Nature},
  volume={400},
  number={6745},
  pages={671--675},
  year={1999},
  publisher={Nature Publishing Group UK London}
}
@article{Eichenbaum1999,
  title={The hippocampus, memory, and place cells: is it spatial memory or a memory space?},
  author={Eichenbaum, Howard and Dudchenko, Paul and Wood, Emma and Shapiro, Matthew and Tanila, Heikki},
  journal={Neuron},
  volume={23},
  number={2},
  pages={209--226},
  year={1999},
  publisher={Elsevier}
}
@article{Nakashiba2009,
  title={Hippocampal CA3 output is crucial for ripple-associated reactivation and consolidation of memory},
  author={Nakashiba, Toshiaki and Buhl, Derek L and McHugh, Thomas J and Tonegawa, Susumu},
  journal={Neuron},
  volume={62},
  number={6},
  pages={781--787},
  year={2009},
  publisher={Elsevier}
}
@article{Ranganath2005,
  title={Functional connectivity with the hippocampus during successful memory formation},
  author={Ranganath, Charan and Heller, Aaron and Cohen, Michael X and Brozinsky, Craig J and Rissman, Jesse},
  journal={Hippocampus},
  volume={15},
  number={8},
  pages={997--1005},
  year={2005},
  publisher={Wiley Online Library}
}
@article{Hamed2016,
  title={Non-parametric analysis of neurochemical effects and Arc expression in amphetamine-induced 50-kHz ultrasonic vocalization},
  author={Hamed, Adam and Daszczuk, Patrycja and Kursa, Miron Bartosz and Turzy{\'n}ska, Danuta and Sobolewska, Alicja and Lehner, Ma{\l}gorzata and Boguszewski, Pawe{\l} M and Szyndler, Janusz},
  journal={Behavioural Brain Research},
  volume={312},
  pages={174--185},
  year={2016},
  publisher={Elsevier}
}
@article{Dudai2004,
  title={The neurobiology of consolidations, or, how stable is the engram?},
  author={Dudai, Yadin},
  journal={Annu. Rev. Psychol.},
  volume={55},
  pages={51--86},
  year={2004},
  publisher={Annual Reviews}
}
@article{Beroun2018,
  title={Generation of silent synapses in dentate gyrus correlates with development of alcohol addiction},
  author={Beroun, Anna and Nalberczak-Sk{\'o}ra, Maria and Harda, Zofia and Piechota, Ma{\l}gorzata and Zi{\'o}{\l}kowska, Magdalena and Ca{\l}y, Anna and Pagano, Roberto and Radwanska, Kasia},
  journal={Neuropsychopharmacology},
  volume={43},
  number={10},
  pages={1989--1999},
  year={2018},
  publisher={Springer International Publishing Cham}
}
@article{Behrens2018,
  title={What is a cognitive map? Organizing knowledge for flexible behavior},
  author={Behrens, Timothy EJ and Muller, Timothy H and Whittington, James CR and Mark, Shirley and Baram, Alon B and Stachenfeld, Kimberly L and Kurth-Nelson, Zeb},
  journal={Neuron},
  volume={100},
  number={2},
  pages={490--509},
  year={2018},
  publisher={Elsevier}
}
@article{Okeefe1978,
  title={Hippocampal place units in the freely moving rat: why they fire where they fire},
  author={O'Keefe, John and Conway, Dulcie H},
  journal={Experimental brain research},
  volume={31},
  pages={573--590},
  year={1978},
  publisher={Springer}
}
@article{Mizumori1999,
  title={Hippocampal representational organization and spatial context},
  author={Mizumori, Sheri JY and Ragozzino, KE and Cooper, BG and Leutgeb, S},
  journal={Hippocampus},
  volume={9},
  number={4},
  pages={444--451},
  year={1999},
  publisher={Wiley Online Library}
}
@article{Dumont2015,
  title={The neural correlates of navigation beyond the hippocampus},
  author={Dumont, Julie R and Taube, Jeffrey S},
  journal={Progress in brain research},
  volume={219},
  pages={83--102},
  year={2015},
  publisher={Elsevier}
}
@article{Grieves2017,
  title={The representation of space in the brain},
  author={Grieves, Roddy M and Jeffery, Kate J},
  journal={Behavioural processes},
  volume={135},
  pages={113--131},
  year={2017},
  publisher={Elsevier}
}
@article{Barry2011,
  title={Imaging spatial learning in the brain using immediate early genes: insights, opportunities and limitations},
  journal={Reviews in Neuroscience},
  author={Barry, Daniel N and Commins, Sean},
  year={2011},
  publisher={Walter de Gruyter}
}
@article{Kubik2007,
  title={Using immediate-early genes to map hippocampal subregional functions},
  author={Kubik, Stepan and Miyashita, Teiko and Guzowski, John F},
  journal={Learning \& memory},
  volume={14},
  number={11},
  pages={758--770},
  year={2007},
  publisher={Cold Spring Harbor Lab}
}
@article{Lisman2005,
  title={The hippocampal-VTA loop: controlling the entry of information into long-term memory},
  author={Lisman, John E and Grace, Anthony A},
  journal={Neuron},
  volume={46},
  number={5},
  pages={703--713},
  year={2005},
  publisher={Elsevier}
}
@article{Hamed2018,
  title={Inter-individual differences in serotonin and glutamate co-transmission reflect differentiation in context-induced conditioned 50-kHz USVs response after morphine withdrawal},
  author={Hamed, Adam and Kursa, Miron Bartosz},
  journal={Brain Structure and Function},
  volume={223},
  pages={3149--3167},
  year={2018},
  publisher={Springer}
}
@article{Bunzeck2011,
  title={Contextual novelty modulates the neural dynamics of reward anticipation},
  author={Bunzeck, Nico and Guitart-Masip, Marc and Dolan, Ray J and D{\"u}zel, Emrah},
  journal={Journal of Neuroscience},
  volume={31},
  number={36},
  pages={12816--12822},
  year={2011},
  publisher={Soc Neuroscience}
}
@article{Hernandez2008,
  title={The hippocampal dentate gyrus is essential for generating contextual memories of fear and drug-induced reward},
  author={Hern{\'a}ndez-Rabaza, V and Hontecillas-Prieto, L and Vel{\'a}zquez-S{\'a}nchez, Clara and Ferragud, A and P{\'e}rez-Villaba, A and Arcusa, A and Barcia, JA and Trejo, Jos{\'e} L and Canales, JJ},
  journal={Neurobiology of learning and memory},
  volume={90},
  number={3},
  pages={553--559},
  year={2008},
  publisher={Elsevier}
}
@article{Czajkowski2014,
  title={Encoding and storage of spatial information in the retrosplenial cortex},
  author={Czajkowski, Rafa{\l} and Jayaprakash, Balaji and Wiltgen, Brian and Rogerson, Thomas and Guzman-Karlsson, Mikael C and Barth, Alison L and Trachtenberg, Joshua T and Silva, Alcino J},
  journal={Proceedings of the National Academy of Sciences},
  volume={111},
  number={23},
  pages={8661--8666},
  year={2014},
  publisher={National Acad Sciences}
}
@article{Milczarek2018,
  title={Spatial memory engram in the mouse retrosplenial cortex},
  author={Milczarek, Michal M and Vann, Seralynne D and Sengpiel, Frank},
  journal={Current Biology},
  volume={28},
  number={12},
  pages={1975--1980},
  year={2018},
  publisher={Elsevier}
}
@article{Alexander2015,
  title={Retrosplenial cortex maps the conjunction of internal and external spaces},
  author={Alexander, Andrew S and Nitz, Douglas A},
  journal={Nature neuroscience},
  volume={18},
  number={8},
  pages={1143--1151},
  year={2015},
  publisher={Nature Publishing Group US New York}
}
@article{Jacob2017,
  title={An independent, landmark-dominated head-direction signal in dysgranular retrosplenial cortex},
  author={Jacob, Pierre-Yves and Casali, Giulio and Spieser, Laure and Page, Hector and Overington, Dorothy and Jeffery, Kate},
  journal={Nature neuroscience},
  volume={20},
  number={2},
  pages={173--175},
  year={2017},
  publisher={Nature Publishing Group US New York}
}
@article{Ikemoto2007,
  title={Dopamine reward circuitry: two projection systems from the ventral midbrain to the nucleus accumbens--olfactory tubercle complex},
  author={Ikemoto, Satoshi},
  journal={Brain research reviews},
  volume={56},
  number={1},
  pages={27--78},
  year={2007},
  publisher={Elsevier}
}
@article{Rossato2009,
  title={Dopamine controls persistence of long-term memory storage},
  author={Rossato, Janine I and Bevilaqua, Lia RM and Izquierdo, Iv{\'a}n and Medina, Jorge H and Cammarota, Mart{\'\i}n},
  journal={Science},
  volume={325},
  number={5943},
  pages={1017--1020},
  year={2009},
  publisher={American Association for the Advancement of Science}
}


@article{mulvihill2018individual,
  title={Individual behavioural predictors of amphetamine-induced emission of 50 kHz vocalization in rats},
  author={Mulvihill, Kevin G and Brudzynski, Stefan M},
  journal={Behavioural brain research},
  volume={350},
  pages={80--86},
  year={2018},
  publisher={Elsevier}
}
@article{taracha2012inter,
  title={Inter-individual diversity and intra-individual stability of amphetamine-induced sensitization of frequency-modulated 50-kHz vocalization in Sprague--Dawley rats},
  author={Taracha, Ewa and Hamed, Adam and Krząścik, Pawe{\l} and Lehner, Ma{\l}gorzata and Sk{\'o}rzewska, Anna and P{\l}a{\'z}nik, Adam and Chrapusta, Stanis{\l}aw J},
  journal={Psychopharmacology},
  volume={222},
  pages={619--632},
  year={2012},
  publisher={Springer}
}

@article{balcerek2021retrosplenial,
  title={Retrosplenial cortex in spatial memory: focus on immediate early genes mapping},
  author={Balcerek, Edyta and W{\l}odkowska, Urszula and Czajkowski, Rafa{\l}},
  journal={Molecular Brain},
  volume={14},
  number={1},
  pages={1--17},
  year={2021},
  publisher={BioMed Central}
}
@article{luo2011linking,
  title={Linking context with reward: a functional circuit from hippocampal CA3 to ventral tegmental area},
  author={Luo, Alice H and Tahsili-Fahadan, Pouya and Wise, Roy A and Lupica, Carl R and Aston-Jones, Gary},
  journal={Science},
  volume={333},
  number={6040},
  pages={353--357},
  year={2011},
  publisher={American Association for the Advancement of Science}
}
@article{sauvage2019single,
  title={Single-cell memory trace imaging with immediate-early genes},
  author={Sauvage, Magdalena and Kitsukawa, Takashi and Atucha, Erika},
  journal={Journal of neuroscience methods},
  volume={326},
  pages={108368},
  year={2019},
  publisher={Elsevier}
}
@article{golebiowska2019serotonin,
  title={Serotonin transporter deficiency alters socioemotional ultrasonic communication in rats},
  author={Golebiowska, Joanna and Ho{\l}uj, Ma{\l}gorzata and Potasiewicz, Agnieszka and Piotrowska, Diana and Kuziak, Agata and Popik, Piotr and Homberg, Judith R and Nikiforuk, Agnieszka},
  journal={Scientific reports},
  volume={9},
  number={1},
  pages={20283},
  year={2019},
  publisher={Nature Publishing Group UK London}
}
@article{lukas2015endogenous,
  title={Endogenous vasopressin, innate anxiety, and the emission of pro-social 50-kHz ultrasonic vocalizations during social play behavior in juvenile rats},
  author={Lukas, Michael and W{\"o}hr, Markus},
  journal={Psychoneuroendocrinology},
  volume={56},
  pages={35--44},
  year={2015},
  publisher={Elsevier}
}
@article{ikemoto1992effects,
  title={The effects of early social isolation on the motivation for social play in juvenile rats},
  author={Ikemoto, Satoshi and Panksepp, Jaak},
  journal={Developmental Psychobiology: The Journal of the International Society for Developmental Psychobiology},
  volume={25},
  number={4},
  pages={261--274},
  year={1992},
  publisher={Wiley Online Library}
}
@article{simola2012pharmacological,
  title={Pharmacological characterization of 50-kHz ultrasonic vocalizations in rats: comparison of the effects of different psychoactive drugs and relevance in drug-induced reward},
  author={Simola, Nicola and Fenu, Sandro and Costa, Giulia and Pinna, Annalisa and Plumitallo, Antonio and Morelli, Micaela},
  journal={Neuropharmacology},
  volume={63},
  number={2},
  pages={224--234},
  year={2012},
  publisher={Elsevier}
}

@article{bijoch2023whole,
  title={Whole-brain tracking of cocaine and sugar rewards processing},
  author={Bijoch, {\L}ukasz and Klos, Joanna and Paw{\l}owska, Monika and Wi{\'s}niewska, Justyna and Legutko, Diana and Szachowicz, Urszula and Kaczmarek, Leszek and Beroun, Anna},
  journal={Translational Psychiatry},
  volume={13},
  number={1},
  pages={20},
  year={2023},
  publisher={Nature Publishing Group UK London}
}
@article{simola2015repeated,
  title={Repeated amphetamine administration and long-term effects on 50-kHz ultrasonic vocalizations: possible relevance to the motivational and dopamine-stimulating properties of the drug},
  author={Simola, Nicola and Morelli, Micaela},
  journal={European Neuropsychopharmacology},
  volume={25},
  number={3},
  pages={343--355},
  year={2015},
  publisher={Elsevier}
}
@article{hamed2016non,
  title={Non-parametric analysis of neurochemical effects and Arc expression in amphetamine-induced 50-kHz ultrasonic vocalization},
  author={Hamed, Adam and Daszczuk, Patrycja and Kursa, Miron Bartosz and Turzy{\'n}ska, Danuta and Sobolewska, Alicja and Lehner, Ma{\l}gorzata and Boguszewski, Pawe{\l} M and Szyndler, Janusz},
  journal={Behavioural Brain Research},
  volume={312},
  pages={174--185},
  year={2016},
  publisher={Elsevier}
}
@article{hamed2020social,
  title={Social deprivation substantially changes multi-structural neurotransmitter signature of social interaction: Glutamate concentration in amygdala and VTA as a key factor in social encounter-induced 50-kHz ultrasonic vocalization},
  author={Hamed, Adam and Kursa, Miron Bartosz},
  journal={European Neuropsychopharmacology},
  volume={37},
  pages={82--99},
  year={2020},
  publisher={Elsevier}
}
@article{hollup2001accumulation,
  title={Accumulation of hippocampal place fields at the goal location in an annular watermaze task},
  author={Hollup, Stig A and Molden, Sturla and Donnett, James G and Moser, May-Britt and Moser, Edvard I},
  journal={Journal of Neuroscience},
  volume={21},
  number={5},
  pages={1635--1644},
  year={2001},
  publisher={Soc Neuroscience}
}
@article{moser2008place,
  title={Place cells, grid cells, and the brain's spatial representation system},
  author={Moser, Edvard I and Kropff, Emilio and Moser, May-Britt},
  journal={Annu. Rev. Neurosci.},
  volume={31},
  pages={69--89},
  year={2008},
  publisher={Annual Reviews}
}
@article{brandon2011reduction,
  title={Reduction of theta rhythm dissociates grid cell spatial periodicity from directional tuning},
  author={Brandon, Mark P and Bogaard, Andrew R and Libby, Christopher P and Connerney, Michael A and Gupta, Kishan and Hasselmo, Michael E},
  journal={Science},
  volume={332},
  number={6029},
  pages={595--599},
  year={2011},
  publisher={American Association for the Advancement of Science}
}
@article{riley1957echolocation,
  title={Echolocation in rats.},
  author={Riley, Donald A and Rosenzweig, Mark R},
  journal={Journal of Comparative and Physiological Psychology},
  volume={50},
  number={4},
  pages={323},
  year={1957},
  publisher={American Psychological Association}
}
@article{rosenzweig1955evidence,
  title={Evidence for echolocation in the rat},
  author={Rosenzweig, Mark R and Riley, Donald A and Krech, David},
  journal={Science},
  volume={121},
  number={3147},
  pages={600--600},
  year={1955},
  publisher={American Association for the Advancement of Science}
}
@article{bell1971echolocation,
  title={Echolocation in the blinded rat},
  author={Bell, Russell A and Noble, ME and Daves, WF},
  journal={Perception \& Psychophysics},
  volume={10},
  pages={112--114},
  year={1971},
  publisher={Springer}
}
@article{geva2015spatial,
  title={Spatial cognition in bats and rats: from sensory acquisition to multiscale maps and navigation},
  author={Geva-Sagiv, Maya and Las, Liora and Yovel, Yossi and Ulanovsky, Nachum},
  journal={Nature Reviews Neuroscience},
  volume={16},
  number={2},
  pages={94--108},
  year={2015},
  publisher={Nature Publishing Group UK London}
}
@article{o1976place,
  title={Place units in the hippocampus of the freely moving rat},
  author={O'Keefe, John},
  journal={Experimental neurology},
  volume={51},
  number={1},
  pages={78--109},
  year={1976},
  publisher={Elsevier}
}
@article{o1971hippocampus,
  title={The hippocampus as a spatial map: preliminary evidence from unit activity in the freely-moving rat.},
  author={O'Keefe, John and Dostrovsky, Jonathan},
  journal={Brain research},
  year={1971},
  publisher={Elsevier Science}
}
@misc{ng2006medical,
  title={Medical image segmentation using k-means clustering and improved watershed algorithm},
  author={Ng, HP and Ong, SH and Foong, KWC and Goh, Poh-Sun and Nowinski, WL},
  booktitle={2006 IEEE southwest symposium on image analysis and interpretation},
  pages={61--65},
  year={2006},
  organization={IEEE}
}
@article{vazdarjanova2004differences,
  title={Differences in hippocampal neuronal population responses to modifications of an environmental context: evidence for distinct, yet complementary, functions of CA3 and CA1 ensembles},
  author={Vazdarjanova, Almira and Guzowski, John F},
  journal={Journal of Neuroscience},
  volume={24},
  number={29},
  pages={6489--6496},
  year={2004},
  publisher={Soc Neuroscience}
}
@article{chawla20043d,
  title={3D-catFISH: a system for automated quantitative three-dimensional compartmental analysis of temporal gene transcription activity imaged by fluorescence in situ hybridization},
  author={Chawla, Monica K and Lin, Gang and Olson, Kathy and Vazdarjanova, Almira and Burke, Sara N and McNaughton, Bruce L and Worley, Paul F and Guzowski, John F and Roysam, Badrinath and Barnes, Carol A},
  journal={Journal of neuroscience methods},
  volume={139},
  number={1},
  pages={13--24},
  year={2004},
  publisher={Elsevier}
}
@article{kubik2012loss,
  title={Loss of activity-dependent Arc gene expression in the retrosplenial cortex after hippocampal inactivation: interaction in a higher-order memory circuit},
  author={Kubik, Stepan and Miyashita, Teiko and Kubik-Zahorodna, Agnieszka and Guzowski, John F},
  journal={Neurobiology of learning and memory},
  volume={97},
  number={1},
  pages={124--131},
  year={2012},
  publisher={Elsevier}
}
@article{mathis2018deeplabcut,
  title={DeepLabCut: markerless pose estimation of user-defined body parts with deep learning},
  author={Mathis, Alexander and Mamidanna, Pranav and Cury, Kevin M and Abe, Taiga and Murthy, Venkatesh N and Mathis, Mackenzie Weygandt and Bethge, Matthias},
  journal={Nature neuroscience},
  volume={21},
  number={9},
  pages={1281--1289},
  year={2018},
  publisher={Nature Publishing Group}
}
@article{jones2000amphetamine,
  title={Amphetamine blocks long-term synaptic depression in the ventral tegmental area},
  author={Jones, Susan and Kornblum, Johanna L and Kauer, Julie A},
  journal={Journal of Neuroscience},
  volume={20},
  number={15},
  pages={5575--5580},
  year={2000},
  publisher={Soc Neuroscience}
}
@article{leith1976amphetamine,
  title={Amphetamine and the reward system: evidence for tolerance and post-drug depression},
  author={Leith, Nancy J and Barrett, Robert J},
  journal={Psychopharmacologia},
  volume={46},
  pages={19--25},
  year={1976},
  publisher={Springer}
}
@article{costa2015involvement,
  title={Involvement of glutamate NMDA receptors in the acute, long-term, and conditioned effects of amphetamine on rat 50kHz ultrasonic vocalizations},
  author={Costa, Giulia and Morelli, Micaela and Simola, Nicola},
  journal={International Journal of Neuropsychopharmacology},
  volume={18},
  number={11},
  year={2015},
  publisher={Oxford University Press}
}
@article{floresco2018differential,
  title={Differential contributions of nucleus accumbens subregions to cue-guided risk/reward decision making and implementation of conditional rules},
  author={Floresco, Stan B and Montes, David R and Maric, MT and van Holstein, Mieke},
  journal={Journal of Neuroscience},
  volume={38},
  number={8},
  pages={1901--1914},
  year={2018},
  publisher={Soc Neuroscience}
}
@article{sohn2022linking,
  title={Linking of NMDA receptors and mGluR5 in the nucleus accumbens core to repeated cocaine-induced 50-kHz ultrasonic vocalization in rats},
  author={Sohn, Sumin and Kim, Sunghyun and Yang, Ju Hwan and Choe, Eun Sang},
  journal={Addiction Biology},
  volume={27},
  number={1},
  pages={e13084},
  year={2022},
  publisher={Wiley Online Library}
}
@article{silkstone2019antagonistic,
  title={The antagonistic relationship between aversive and appetitive emotional states in rats as studied by pharmacologically-induced ultrasonic vocalization from the nucleus accumbens and lateral septum},
  author={Silkstone, Michael and Brudzynski, Stefan M},
  journal={Pharmacology Biochemistry and Behavior},
  volume={181},
  pages={77--85},
  year={2019},
  publisher={Elsevier}
}

@article{hori2013tickling,
  title={Tickling increases dopamine release in the nucleus accumbens and 50 kHz ultrasonic vocalizations in adolescent rats},
  author={Hori, Miyo and Shimoju, Rie and Tokunaga, Ryota and Ohkubo, Masato and Miyabe, Shigeki and Ohnishi, Junji and Murakami, Kazuo and Kurosawa, Mieko},
  journal={Neuroreport},
  volume={24},
  number={5},
  pages={241--245},
  year={2013},
  publisher={LWW}
}
@article{brudzynski2012quinpirole,
  title={Quinpirole-induced 50 kHz ultrasonic vocalization in the rat: role of D2 and D3 dopamine receptors},
  author={Brudzynski, Stefan M and Komadoski, Melanie and Pierre, Joel St},
  journal={Behavioural Brain Research},
  volume={226},
  number={2},
  pages={511--518},
  year={2012},
  publisher={Elsevier}
}

@article{burgdorf2001nucleus,
  title={Nucleus accumbens amphetamine microinjections unconditionally elicit 50-kHz ultrasonic vocalizations in rats.},
  author={Burgdorf, Jeffrey and Knutson, Brian and Panksepp, Jaak and Ikemoto, Satoshi},
  journal={Behavioral neuroscience},
  volume={115},
  number={4},
  pages={940},
  year={2001},
  publisher={American Psychological Association}
}
@article{costa2020influence,
  title={Influence of dopamine transmission in the medial prefrontal cortex and dorsal striatum on the emission of 50-kHz ultrasonic vocalizations in rats treated with amphetamine: Effects on drug-stimulated and conditioned calls},
  author={Costa, Giulia and Serra, Marcello and Marongiu, Jacopo and Morelli, Micaela and Simola, Nicola},
  journal={Progress in Neuro-Psychopharmacology and Biological Psychiatry},
  volume={97},
  pages={109797},
  year={2020},
  publisher={Elsevier}
}
@article{guo2013dbs,
  title={DBS of nucleus accumbens on heroin seeking behaviors in self-administering rats},
  author={Guo, Liemei and Zhou, Hongyu and Wang, Ran and Xu, Jiwen and Zhou, Wenhua and Zhang, Fuqiang and Tang, Shuaien and Liu, Huifen and Jiang, Jiyao},
  journal={Drug and alcohol dependence},
  volume={129},
  number={1-2},
  pages={70--81},
  year={2013},
  publisher={Elsevier}
}
@article{willuhn2014phasic,
  title={Phasic dopamine release in the nucleus accumbens in response to pro-social 50 kHz ultrasonic vocalizations in rats},
  author={Willuhn, Ingo and Tose, Amanda and Wanat, Matthew J and Hart, Andrew S and Hollon, Nick G and Phillips, Paul EM and Schwarting, Rainer KW and W{\"o}hr, Markus},
  journal={Journal of Neuroscience},
  volume={34},
  number={32},
  pages={10616--10623},
  year={2014},
  publisher={Soc Neuroscience}
}
@article{humphries2010ventral,
  title={The ventral basal ganglia, a selection mechanism at the crossroads of space, strategy, and reward.},
  author={Humphries, Mark D and Prescott, Tony J},
  journal={Progress in neurobiology},
  volume={90},
  number={4},
  pages={385--417},
  year={2010},
  publisher={Elsevier}
}
@article{bornstein2011multiplicity,
  title={Multiplicity of control in the basal ganglia: computational roles of striatal subregions},
  author={Bornstein, Aaron M and Daw, Nathaniel D},
  journal={Current opinion in neurobiology},
  volume={21},
  number={3},
  pages={374--380},
  year={2011},
  publisher={Elsevier}
}
@article{mannella2013nucleus,
  title={The nucleus accumbens as a nexus between values and goals in goal-directed behavior: a review and a new hypothesis},
  author={Mannella, Francesco and Gurney, Kevin and Baldassarre, Gianluca},
  journal={Frontiers in behavioral neuroscience},
  volume={7},
  pages={135},
  year={2013},
  publisher={Frontiers Media SA}
}
@article{penner2012neural,
  title={Neural systems analysis of decision making during goal-directed navigation},
  author={Penner, Marsha R and Mizumori, Sheri JY},
  journal={Progress in neurobiology},
  volume={96},
  number={1},
  pages={96--135},
  year={2012},
  publisher={Elsevier}
}
@article{martin2000effects,
  title={Effects of reward anticipation, reward presentation, and spatial parameters on the firing of single neurons recorded in the subiculum and nucleus accumbens of freely moving rats},
  author={Martin, Patrick D and Ono, Taketoshi},
  journal={Behavioural brain research},
  volume={116},
  number={1},
  pages={23--38},
  year={2000},
  publisher={Elsevier}
}
@article{khamassi2012integrating,
  title={Integrating cortico-limbic-basal ganglia architectures for learning model-based and model-free navigation strategies},
  author={Khamassi, Mehdi and Humphries, Mark D},
  journal={Frontiers in behavioral neuroscience},
  volume={6},
  pages={79},
  year={2012},
  publisher={Frontiers Media SA}
}
@article{groenewegen1982cortical,
  title={Cortical afferents of the nucleus accumbens in the cat, studied with anterograde and retrograde transport techniques},
  author={Groenewegen, HJ and Room, P and Witter, MP and Lohman, AHM},
  journal={Neuroscience},
  volume={7},
  number={4},
  pages={977--996},
  year={1982},
  publisher={Elsevier}
}
@article{jayaraman1985organization,
  title={Organization of thalamic projections in the nucleus accumbens and the caudate nucleus in cats and its relation with hippocampal and other subcortical afferents},
  author={Jayaraman, A},
  journal={Journal of Comparative Neurology},
  volume={231},
  number={3},
  pages={396--420},
  year={1985},
  publisher={Wiley Online Library}
}
@article{powell1976connections,
  title={Connections of the nucleus accumbens},
  author={Powell, Ervin W and Leman, Robert B},
  journal={Brain research},
  volume={105},
  number={3},
  pages={389--403},
  year={1976},
  publisher={Elsevier}
}
@article{zahm1992significance,
  title={On the significance of subterritories in the “accumbens” part of the rat ventral striatum},
  author={Zahm, DS and Brog, JS},
  journal={Neuroscience},
  volume={50},
  number={4},
  pages={751--767},
  year={1992},
  publisher={Elsevier}
}
@article{jin2022examining,
  title={Examining ventral subiculum and basolateral amygdala projections to the nucleus accumbens shell: Differential expression of VGLuT1, VGLuT2 and VGaT in the rat},
  author={Jin, Shubo and Maddern, Xavier J and Campbell, Erin J and Lawrence, Andrew J},
  journal={Neuroscience Letters},
  volume={788},
  pages={136858},
  year={2022},
  publisher={Elsevier}
}
@article{van2020basolateral,
  title={Basolateral amygdala--nucleus accumbens circuitry regulates optimal cue-guided risk/reward decision making},
  author={van Holstein, Mieke and MacLeod, Paula E and Floresco, Stan B},
  journal={Progress in Neuro-Psychopharmacology and Biological Psychiatry},
  volume={98},
  pages={109830},
  year={2020},
  publisher={Elsevier}
}
@article{zahm1998caudomedial,
  title={Is the caudomedial shell of the nucleus accumbens part of the extended amygdala? A consideration of connections},
  author={Zahm, Daniel S},
  journal={Critical Reviews™ in Neurobiology},
  volume={12},
  number={3},
  year={1998},
  publisher={Begel House Inc.}
}
@article{yim1982response,
  title={Response of nucleus accumbens neurons to amygdala stimulation and its modification by dopamine},
  author={Yim, Chi Yiu and Mogenson, Gordon J},
  journal={Brain research},
  volume={239},
  number={2},
  pages={401--415},
  year={1982},
  publisher={Elsevier}
}
@article{di1999drug,
  title={Drug addiction as a disorder of associative learning: role of nucleus accumbens shell/extended amygdala dopamine},
  author={DI CHIARA, GAETANO and Tanda, G and BASSAREO, VALENTINA and Pontieri, F and Acquas, ELIO and FENU, SANDRO and Cadoni, C and CARBONI, EZIO},
  journal={Annals of the New York Academy of Sciences},
  volume={877},
  number={1},
  pages={461--485},
  year={1999},
  publisher={Wiley Online Library}
}
@article{yu2022contingent,
  title={Contingent Amygdala Inputs Trigger Heterosynaptic LTP at Hippocampus-To-Accumbens Synapses},
  author={Yu, Jun and Sesack, Susan R and Huang, Yanhua and Schl{\"u}ter, Oliver M and Grace, Anthony A and Dong, Yan},
  journal={Journal of Neuroscience},
  volume={42},
  number={34},
  pages={6581--6592},
  year={2022},
  publisher={Soc Neuroscience}
}
@article{martin2001locomotion,
  title={Locomotion towards a goal alters the synchronous firing of neurons recorded simultaneously in the subiculum and nucleus accumbens of rats},
  author={Martin, Patrick D},
  journal={Behavioural brain research},
  volume={124},
  number={1},
  pages={19--28},
  year={2001},
  publisher={Elsevier}
}
@article{friedman2002comparison,
  title={Comparison of hippocampal, amygdala, and perirhinal projections to the nucleus accumbens: combined anterograde and retrograde tracing study in the Macaque brain},
  author={Friedman, David P and Aggleton, John P and Saunders, Richard C},
  journal={Journal of Comparative Neurology},
  volume={450},
  number={4},
  pages={345--365},
  year={2002},
  publisher={Wiley Online Library}
}
@article{reynolds2008emotional,
  title={Emotional environments retune the valence of appetitive versus fearful functions in nucleus accumbens},
  author={Reynolds, Sheila M and Berridge, Kent C},
  journal={Nature neuroscience},
  volume={11},
  number={4},
  pages={423--425},
  year={2008},
  publisher={Nature Publishing Group US New York}
}
@article{yang2018nucleus,
  title={Nucleus accumbens subnuclei regulate motivated behavior via direct inhibition and disinhibition of VTA dopamine subpopulations},
  author={Yang, Hongbin and de Jong, Johannes W and Tak, YeEun and Peck, James and Bateup, Helen S and Lammel, Stephan},
  journal={Neuron},
  volume={97},
  number={2},
  pages={434--449},
  year={2018},
  publisher={Elsevier}
}
@article{arias2010dopaminergic,
  title={Dopaminergic reward system: a short integrative review},
  author={Arias-Carri{\'o}n, Oscar and Stamelou, Maria and Murillo-Rodr{\'\i}guez, Eric and Men{\'e}ndez-Gonz{\'a}lez, Manuel and P{\"o}ppel, Ernst},
  journal={International archives of medicine},
  volume={3},
  number={1},
  pages={1--6},
  year={2010},
  publisher={BioMed Central}
}
@article{polter2014stress,
  title={Stress and VTA synapses: implications for addiction and depression},
  author={Polter, Abigail M and Kauer, Julie A},
  journal={European journal of neuroscience},
  volume={39},
  number={7},
  pages={1179--1188},
  year={2014},
  publisher={Wiley Online Library}
}
@article{cohen2022reward,
  title={Reward enhances memory via age-varying online and offline neural mechanisms across development},
  author={Cohen, Alexandra O and Glover, Morgan M and Shen, Xinxu and Phaneuf, Camille V and Avallone, Kristen N and Davachi, Lila and Hartley, Catherine A},
  journal={Journal of Neuroscience},
  volume={42},
  number={33},
  pages={6424--6434},
  year={2022},
  publisher={Soc Neuroscience}
}
@article{diaz2021bidirectional,
  title={Bidirectional role of dopamine in learning and memory-active forgetting},
  author={D{\'\i}az, Fernando Castillo and Caffino, Lucia and Fumagalli, Fabio},
  journal={Neuroscience \& Biobehavioral Reviews},
  volume={131},
  pages={953--963},
  year={2021},
  publisher={Elsevier}
}
@article{hainmueller2020dentate,
  title={Dentate gyrus circuits for encoding, retrieval and discrimination of episodic memories},
  author={Hainmueller, Thomas and Bartos, Marlene},
  journal={Nature Reviews Neuroscience},
  volume={21},
  number={3},
  pages={153--168},
  year={2020},
  publisher={Nature Publishing Group UK London}
}
@article{kursa2010feature,
  title={Feature selection with the Boruta package},
  author={Kursa, Miron B and Rudnicki, Witold R},
  journal={Journal of statistical software},
  volume={36},
  pages={1--13},
  year={2010}
}
@article{holm1979simple,
  title={A simple sequentially rejective multiple test procedure},
  author={Holm, Sture},
  journal={Scandinavian journal of statistics},
  pages={65--70},
  year={1979},
  publisher={JSTOR}
}
@article{jaworski2018c,
  title={c-Fos and neuronal plasticity: the aftermath of Kaczmarek’s theory},
  author={Jaworski, Jacek and Kalita, Katarzyna and Knapska, Ewelina},
  journal={Acta Neurobiol Exp (Wars)},
  volume={78},
  number={4},
  pages={287--296},
  year={2018}
}
@article{guzowski2001cellular,
  title={Cellular compartment analysis of temporal activity by fluorescence in situ hybridization (catFISH): current protocols in neuroscience/editorial board, Jacqueline N. Crawley [et al]},
  author={Guzowski, JF and Worley, PF},
  journal={Chapter},
  volume={1},
  year={2001}
}
@book{paxinos2006rat,
  title={The rat brain in stereotaxic coordinates: hard cover edition},
  author={Paxinos, George and Watson, Charles},
  year={2006},
  publisher={Elsevier}
}
