## Supplementary material for "Spatio-temporal mechanisms of consolidation, recall and reconsolidation in reward-related memory trace": Manuscript LaTeX source: main.pdf

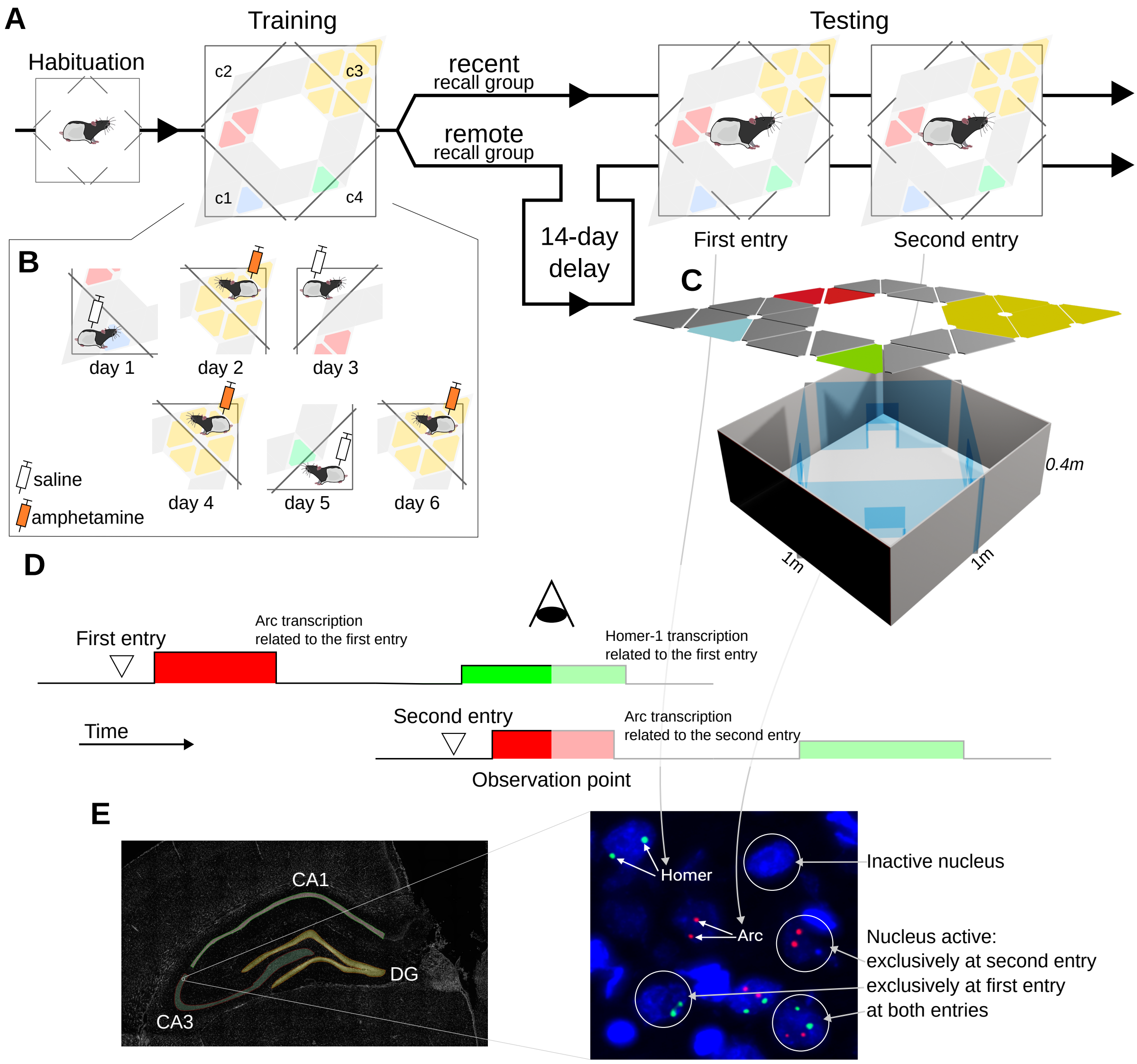


**Fig. 1** Overview of the experimental procedure. A. Each rat is habituated to an experimental cage, trained to associate corner c3 with amphetamine effects and undergoes testing which consists of two entries into the context. Rats are divided into recent and remote recall groups, which differ only in an additional two-week delay between training and testing sessions in the remote group. B. Details of the 6-day training routine; on even days rat is placed in the conditioned corner c2 and given amphetamine, while on odd days placed in each other corner and given sham, saline injection. C. Scheme of the experimental cage; a $1m \times 1m$ box is partitioned into 5 regions by transparent walls equipped with doors that can be opened (during habituation and testing) or closed (during training). An over-hanged illumination assembly provides animals with spatial cues. D. Scheme of the CatFISH method used to elucidate changes in brain activity between test entries. mRNAs of two IEGs, Arc and Homer, are detected; the temporal distribution of entries is set up so that at the observation point, we can independently detect expression of IEGs triggered by the activity at either test entry. E. Image analysis pipeline; brain structures are identified and marked as ROIs using shapes from a reference atlas [33]. Within each ROI, we identify nuclei and, according to IEG expression detected within them, classify into inactive, active during eith erentry and active during both entries.

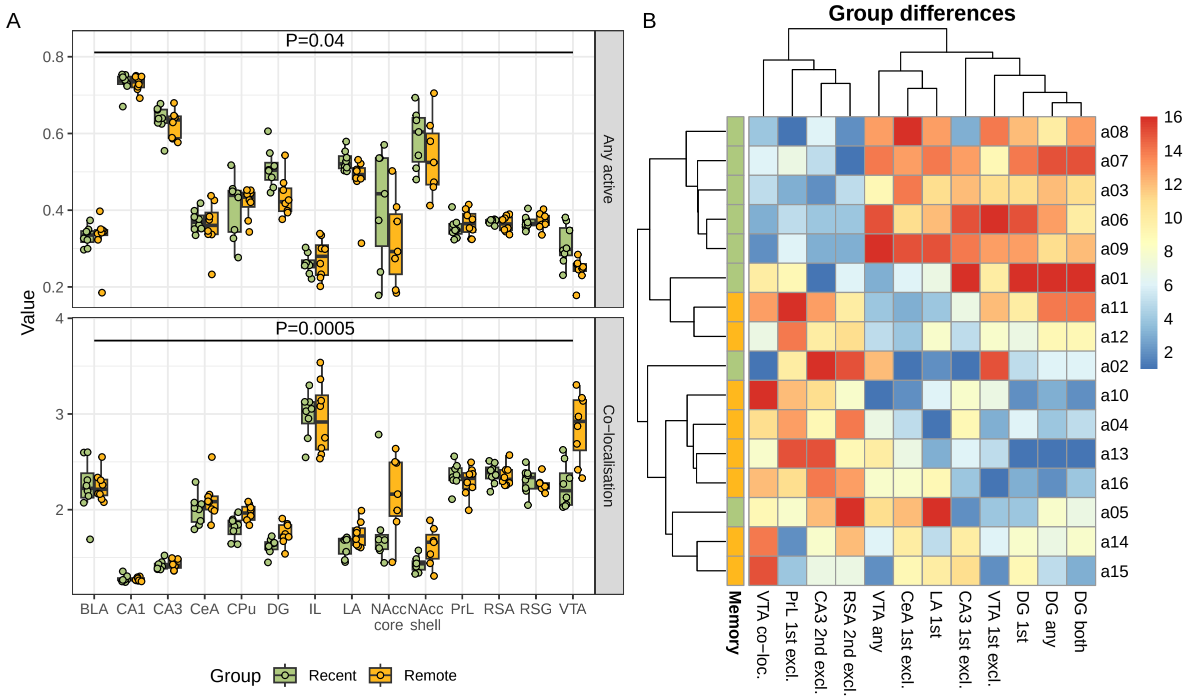


**Fig. 2** The differences in overall activity and activity co-localisation in investigated structures between rats with recent and remote recall. A. Comparison of the fraction of active nuclei (top) and co-localisation coefficient (bottom); both conditioned on the brain structure. In the remote recall, the activity has generally dropped in comparison to the recent recall, while the co-localisation has increased, which suggests the development of specialised neuron sub-populations. Box-plots adhere to a standard Tukey's definition, original data is superimposed as jittered points. B. Heatmap showing the values of brain activity descriptors found to significantly differ between groups in a machine learning-based multivariate analysis. Most straightforward effects were identified in the VTA and hippocampus, but also in cortical regions and amygdala. Values were shown as ranks for clarity, with rank one given to a lowest value (blue) and rank sixteen to the highest (red).

#### 2.4 High, memory remoteness-independent variation in NAcc is explained by behaviour

As mentioned in Section 2.2, there is a substantial variation of activity in certain structures which is not explained by the recent/remote group; in particular in the nucleus accumbens. By using the trajectory data extracted from the video recordings of the behaviour, we have quantified the inclination of a rat to stay in the corner that was previously associated with the amphetamine injection. To rule out the impact of variable exploration tendencies, we have expressed it as a correctness score, defined as a fraction of the total time spent in any corner which was spent in a conditioned one.

While this score was not significantly correlated with the recent/remote recall group, we have identified significant molecular associations with machine learning; they are collected in Figure 3.

High correctness was identified to be explained by the activity of hippocampal CA3 as well as both the shell and core of the nucleus accumbens. Moreover, these activity levels were fairly consistent across two entries, henceforth almost all pattern parameters were selected. A similar pattern applies to caudate putamen (CPu), yet to a lesser extent, because CPu activity at the second entry is less discriminating.

Finally, there are also significant correlations between first entry correctness and overall activity of the lateral amygdala (LA) as well as activity co-localisation in the granular retrosplenial cortex (RSG).

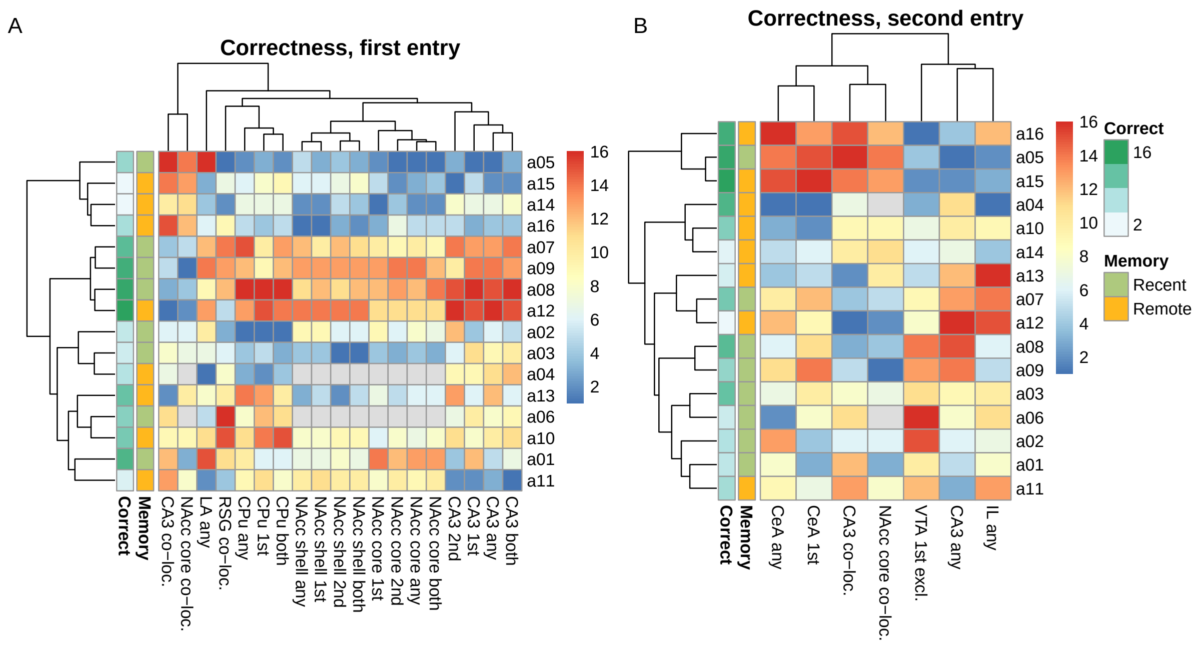


**Fig. 3** The factors associated with the correctness in the first test entry (A) and second test entry (B). Heatmaps show the values of brain activity descriptors found to be significantly associated with correctness in a machine learning-based multivariate analysis. The high first-entry correctness score was related to nucleus accumbens and CA3 activity, while the central amygdala was a key driver in the second entry. Values were shown as ranks for clarity, with rank one given to a lowest value (blue) and rank sixteen to the highest (red); gray encodes missing values.

#### 2.5 Behaviour at each entry has distinct neuronal activity correlates

While the overall activity in NAcc and CA3, which are strong correlates of correctness at the first entry, remains fairly consistent between entries, they cease to strongly explain the correctness at the second entry. Only co-localisations at CA3 and NAcc core and overall activity of CA3 remain to be important.

On the other hand, the activity of the central amygdala (CeA) becomes a key driver, we also see significant interactions involving the infralimbic cortex (IL) and VTA. They are rather complex and involve a structure state at the first entry.

#### 2.6 Strongest inter-structural co-activation occurs at second entry

We have investigated activity correlations across all investigated neuronal sub-populations in all structures. The graphs constructed from these identified to be statistically significant ones are presented in Figure 4 — there were none such in the *Any* and *Second* classes.

This analysis has mostly uncovered trivial, inter-structural correlations, in particular within amygdala (BLA-LA-CeA), NAcc (shell & core), retrosplenial cortex (RSG-RSA) and hippocampus (CA1-CA3, though not DG).

When analysing populations active exclusively during the first entry, we have found a significant negative correlation between CeA and PrL. In comparison to the general first entry activity graphs, we see that both the internal coupling in NAcc and between CA1-CA3 have disappeared, which indicates that this synchronisation has persisted over both entries. On the other hand, RSG-RSA link is present in both exclusive views but not in any more general one, suggesting it is a result of a more temporally localised phenomenon.

Most of the significant correlations identified are between the sizes of neuronal populations active exclusively during the second entry. Here, we see intra-structural coherence in the amygdala,

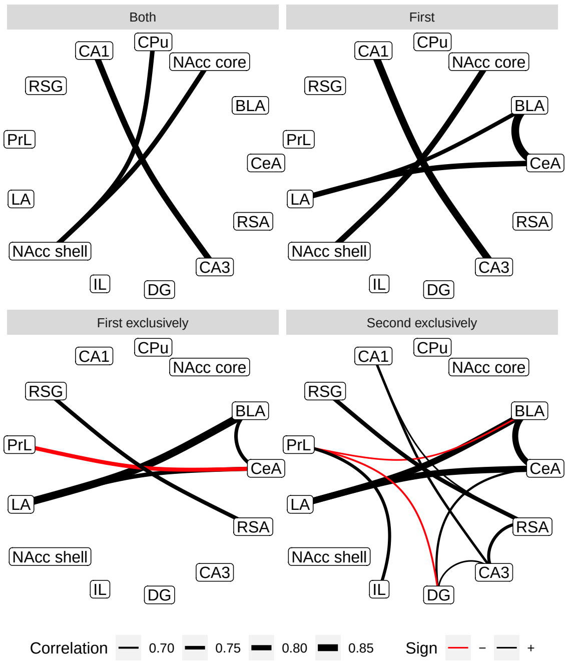


**Fig. 4** Significant monotonic correlations between values of activity descriptors between investigated brain structures. Each panel presents graph for a certain activity descriptor; panels with no links (for an activity at any entry, *any*, and at the second entry, *2nd*) were omitted. The densest, non-trivial networks can be observed between sizes of neuron sub-populations active exclusively at certain entry. The weight of a link corresponds to an absolute value of Spearman correlation coefficient; black links represent positive, while red ones negative correlations.

### Funding

R.C., M.F. and A.H. were supported by grant Sonata-bis 2014/14/E/NZ4/00172 from the National Science Centre Poland. A.H. and W.K. were supported by grant Opus 2018/29/B/NZ7/02021 from the National Science Centre Poland.

### Acknowledgments

We wish to thank Tomasz Gomoła for the construction a 3D model of an experimental setup.

### 5 Methods

#### 5.1 Animals

Adult male Long-Evans rats ($n = 18$, $180 \pm 20g$) were used in the experiment. Animals were purchased from a licensed breeder (the Polish Academy of Science Medical Research Center, Warsaw, Poland), and housed in standard laboratory conditions under $12h$:$12h$ light:dark cycles (lights on at 7 a.m.), at a constant temperature ($21 \pm 2°C$) with 70% humidity. Rats had free access to both food and water. All experiments were performed in accordance with the European Communities Council Directive of 24 November 1986 (86/609 EEC). Local Ethical Committee approved all experimental procedures using animal subjects (539/2018).

#### 5.2 Behavioural experiment

All experiments utilised the same $1m \times 1m$ cage, partitioned into 5 regions (four corners and a central interconnecting space) with translucent walls equipped with doors. In the *training* configuration doors are closed, henceforth a rat can be confined in a selected corner but retains the capacity to observe the whole cage through the wall. On

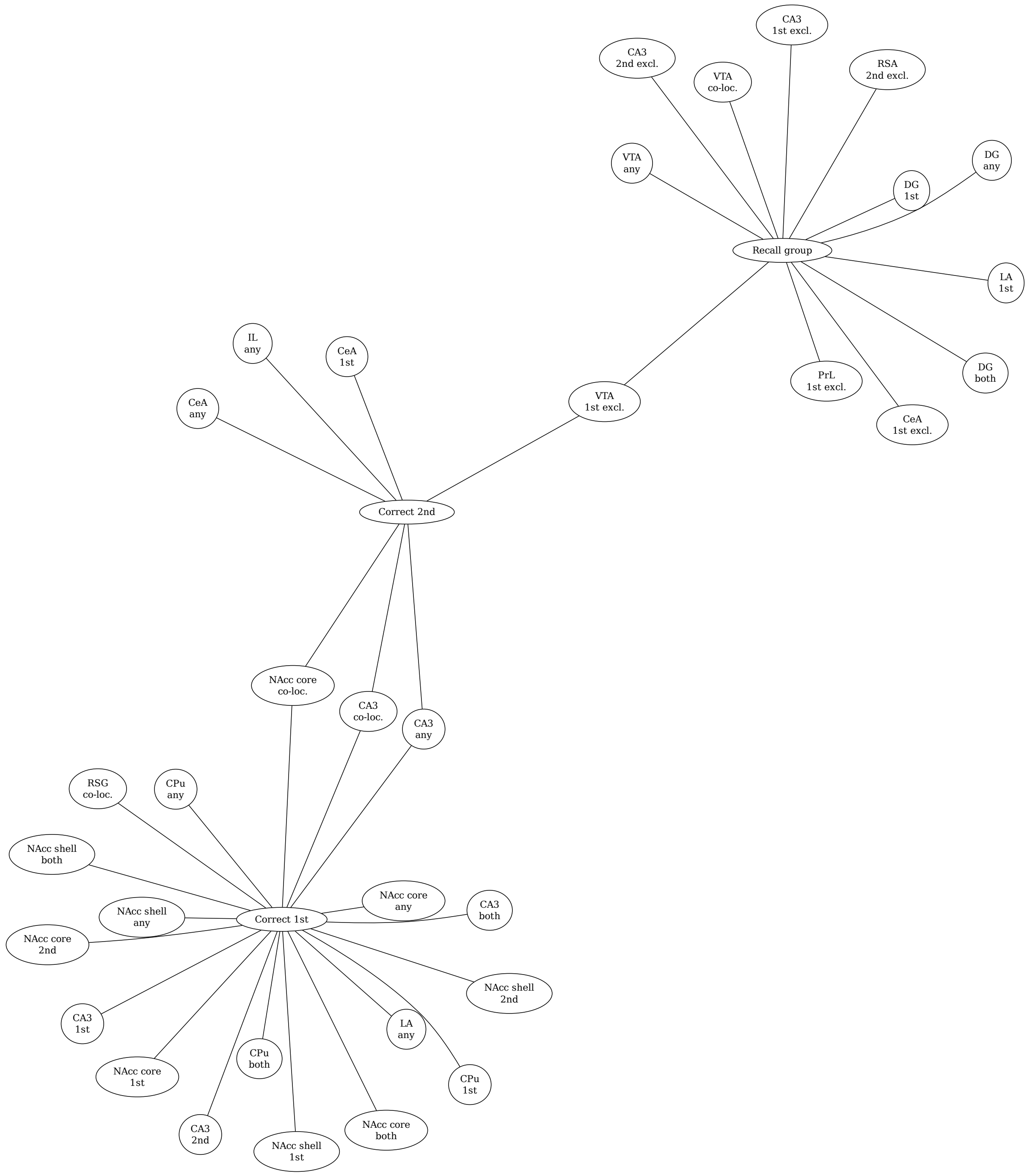


**Fig. 5** Overview of the overlap between machine learning-selected significantly important factors connected with the memory recall group and correctness score in either test entry. Although hippocampus, amygdala and nucleus accumbens re-appear in all contexts, their involvement differs at the level of finer-grained neuronal populations.

#### 5.4 Image acquisition and analysis

Stained slices containing representative samples of interesting structures were scanned using Inverted Leica DMI 6000 microscope with Andor DSD2 Confocal Module and HC PL APO CS2 20x/0.70 Immersive objective. Fluorescence was excited with 200W halogen lamp. To detect nuclei, Homer-1a and Arc, three fluorescent filtersets were used, respectively: DAPI (exc. $390\pm40nm$, 405 DM, em. $452\pm45nm$), FITC (exc. $483\pm28nm$, 488 DM, em. $525\pm45nm$) and Cy3 (exc. $556\pm20nm$, 561 DM, em. $609\pm54nm$).
