## Supplementary material for "Spatio-temporal mechanisms of consolidation, recall and reconsolidation in reward-related memory trace": Manuscript LaTeX source: main.tex

\documentclass[sn-nature,iicol,pdflatex]{sn-jnl}

\usepackage[utf8]{inputenc}
\usepackage[T1]{fontenc}
\usepackage{graphicx}
\usepackage{multirow}%
\usepackage{amsmath,amssymb,amsfonts}%
\usepackage{amsthm}%
\usepackage{mathrsfs}%
\usepackage[title]{appendix}%
\usepackage{xcolor}%
\usepackage{textcomp}%
\usepackage{manyfoot}%
\usepackage{booktabs}%
\usepackage{algorithm}%
\usepackage{algorithmicx}%
\usepackage{algpseudocode}%
\usepackage{listings}%
\usepackage{lineno}
\usepackage{gensymb}
\hyphenation{ac-cum-bens}

\begin{document}
 \title{Spatio-temporal mechanisms of consolidation, recall and reconsolidation in reward-related memory trace}                               
\author*[1]{Adam Hamed}
\author[2]{Miron Bartosz Kursa}
\author[1]{Wiktoria Karwicka}
\author[2]{Krzysztof Piotr Piwoński}
\author[1]{Monika Falińska}
\author[3]{Konrad Danielewski}
\author[4]{Emilia Rejmak-Kozicka}
\author[1]{Urszula Włodkowska}
\author[5]{Stepan Kubik}
\author*[1]{Rafał Czajkowski}
\affil[1]{Laboratory of Spatial Memory, Nencki Institute of Experimental Biology, Polish Academy of Sciences}
\affil[2]{Interdisciplinary Centre for Mathematical and Computational Modelling, University of Warsaw}
\affil[3]{Laboratory of Emotions Neurobiology, Nencki Institute of Experimental Biology, Polish Academy of Sciences}
\affil[4]{BRAINCITY, Nencki Institute of Experimental Biology, Polish Academy of Sciences}
\affil[5]{Institute of Physiology, Academy of Sciences of the Czech Republic}
\abstract{The formation of memories is a complex, multi-scale phenomenon, especially when it involves integration of information from various brain systems.
We have investigated the differences between a novel and consolidated association of spatial cues and amphetamine administration, using an in-situ hybridisation method to track the short-term dynamics during the recall testing. 
We have found that remote recall group involves smaller, but more consolidated groups of neurons, which is consistent with their specialisation.
By employing machine learning analysis, we have shown this pattern is especially pronounced in the VTA; furthermore, we also uncovered significant activity patterns in retrosplenial and prefrontal cortices, as well as in the DG and CA3 subfields of the hippocampus.
The behavioural propensity towards the associated localisation appears to be driven by the nucleus accumbens, however, further modulated by a trio of the amygdala, VTA and hippocampus, as the trained association is confronted with test experience.
These results show that memory mechanisms must be modelled considering individual differences in motivation, as well as covering dynamics of the process.}

\maketitle

\section{Introduction}


Preserving the memory of an important life event requires successful integration of several modalities that constitute a multisensory trace, often referred to as the memory engram \cite{Dudai2004}.
In particular, spatial memory is defined as the brain’s ability to encode key features of the external environment and to navigate within the boundaries of this mental representation, also known as cognitive map \cite{Behrens2018,Okeefe1978}.
On the physiological level, it manifests as populations of neurons exhibiting activity tuned to specific aspects of the external spatial context, in particular, firing correlated with an animal's presence in a certain, unique location within the environment \cite{Mizumori1999,hollup2001accumulation}. 
Such neuron populations, called place cells \cite{o1971hippocampus, o1976place}, are mainly located in the hippocampus, yet their specificity is influenced by several other groups of spatially tuned neurons, most notably grid cells, border cells and head direction cells \cite{moser2008place, Dumont2015, Grieves2017,brandon2011reduction}.
These cells are in turn distributed among several cortical and sub-cortical structures.
Said network is further connected by a plethora of direct and indirect projections, serving both top-down and bottom-up processing of spatial information and associating it with representations stored by other circuits.

In the case of episodic memories, their spatial context can be, among others, linked to the emotional state during learning.
This phenomenon is often investigated with aversive states, like those triggered by physical pain or alarming stimuli.
However, it occurs regardless of the valence, and can also involve appetitive states; in particular reward-related ones, on which we focus in this work.
These are processed by yet another crucial functional element of the brain: the reward system, responsible for cognitive fundamentals of motivation and reinforcement, as well as a crucial agent in many psychological disorders.
The physiological connections between these systems have also been established; for instance, it was demonstrated that a projection from the hippocampus to the ventral tegmental area (VTA) mediated relations between context and reward \cite{Lisman2005,luo2011linking}.

Appetitive states can be naturally evoked by likes of attractive foods or positive social interactions \cite{golebiowska2019serotonin, ikemoto1992effects,hamed2020social}, 
but also induced with substances that directly activate the reward system, that is pharmacological rewards \cite{simola2012pharmacological, hamed2016non, bijoch2023whole}. 
One of them is an amphetamine, which is known to reliably activate the reward system \cite{leith1976amphetamine,jones2000amphetamine,mulvihill2018individual} and to induce the emission of 50kHz band ultrasonic vocalisations, which are a marker of affective appetitive states in rats \cite{costa2015involvement,taracha2012inter, Hamed2016}.
To this end, it is often used to induce a consistent appetitive emotional state to be linked with spatial cues during the experiment.

The immense complexity of brain mechanics makes their quantification a highly non-trivial endeavour.
Henceforth, contemporary approaches have to focus on narrow aspects of activity, especially those accessible for measurements and relatively straightforward to interpret.
On the transcriptomic level, one of such convenient manifestations is the transient expression of immediate early genes (IEGs), in particular, cFos, Arc, Homer, and zif268 \cite{Barry2011,balcerek2021retrosplenial,jaworski2018c}. 
They can be used to visualise and quantify the cellular ensembles involved in the memory trace, as demonstrated in \cite{Kubik2007, sauvage2019single}. 

The memory has a very rich dynamic \cite{Squire2015}.
In a global view, it consists of memory formation, consolidation, recognition, recall,  re-consolidation and extinction; yet all of these processes have their own local dynamics of variable time scales and involve complex phenomena, for instance, engram shifts between brain structures. 
Consequently, the investigation of temporal aspects is crucial for the analyses of memory mechanics.
Thanks to the previously identified distinctive patterns of IEG expression in time, they are an important tool for this task.

In this work, we aim to investigate the neural mechanisms involved in the interplay of spatial memory and reward processing systems, underlying the emotional perception of context, which is in turn crucial to understanding goal-directed behaviour associated with reward-seeking, as well as spatial memory storage, maturation and consolidation.
Rodent models offer a way to explore these relationships experimentally on the behavioural and molecular levels. 
One of the simplest and most commonly used models is the \emph{conditioned place preference} (CPP) paradigm, where the animal learns to associate a particular area of experimental enclosure (facilitated by spatial cues) with an emotional experience.
We have devised an approach extending CPP with an open-field paradigm, orchestrating a reward-seeking task based on spatial cues in a 5-region cage (with 4 corners and interconnecting centre space).
For conditioning, we have used amphetamine administration in one of the corners.

To investigate the mechanism underlying such evoked memory, we have used the 
CatFISH method (Cellular compartment analysis of temporal activity by fluorescent in situ hybridization) to measure the co-expression of two IEGs, Arc and Homer-1A, in nine brain areas covering both the cortex and sub-cortical structures \cite{guzowski2001cellular}. 
In order to elucidate larger-scale temporal dynamics, we have analysed two groups of rats, with a recent and remote memory recall.
By using machine learning, we were able to pinpoint the circuits responsible for behavioural responses related to reward-seeking based on spatial cues.


\section{Results}
\subsection{Experiment overview}
Rats were trained to associate a particular corner of a rectangular enclosure with the effects of amphetamine injection.
In training, doors in cage partitions were closed, constraining rats to stay in a corner they were placed. 
On the other hand, for testing, said doors were opened, and rats were allowed to freely roam around the cage; yet, they were not given amphetamine.

The testing session included two, five-minute entries, separated by a 20-minute break; afterwards, rats were sacrificed and their brains analysed (Figure~\ref{fig:scheme}).
Due to such a course, we were able to exploit the temporal expression patterns of two IEGs to untangle the brain activity in either entry.
Precisely, we were able to measure what fraction of neuronal nuclei within a given brain structure was active during any entry, either entry, exclusively either entry and finally during both entries (Figure~\ref{fig:scheme} E).
Out of these descriptors, we have also calculated the co-localisation coefficient, expressing the degree to which consistently the same neurons exhibit activity across entries.

\begin{figure*}
 \centering
 \includegraphics[width=\textwidth]{scheme.pdf}
 \caption{\label{fig:scheme} Overview of the experimental procedure. 
A. Each rat is habituated to an experimental cage, trained to associate corner c3 with amphetamine effects and undergoes testing which consists of two entries into the context.
Rats are divided into recent and remote recall groups, which differ only in an additional two-week delay between training and testing sessions in the remote group.
B. Details of the 6-day training routine; on even days rat is placed in the conditioned corner c2 and given amphetamine, while on odd days placed in each other corner and given sham, saline injection.
C. Scheme of the experimental cage; a 1$m\times 1m$ box is partitioned into 5 regions by transparent walls equipped with doors that can be opened (during habituation and testing) or closed (during training). 
An over-hanged illumination assembly provides animals with spatial cues.
D. Scheme of the CatFISH method used to elucidate changes in brain activity between test entries.
mRNAs of two IEGs, Arc and Homer, are detected; the temporal distribution of entries is set up so that at the observation point, we can independently detect expression of IEGs triggered by the activity at either test entry.
E. Image analysis pipeline; brain structures are identified and marked as ROIs using shapes from a reference atlas \cite{paxinos2006rat}.
Within each ROI, we identify nuclei and, according to IEG expression detected within them, classify into inactive, active during eith erentry and active during both entries.}
\end{figure*}

This experiment was repeated in two variants, investigating recent and remote (consolidated) recall; the first one was carried out the day after training, while the second one was two weeks after the last training session.
Both sessions were video-recorded, which allowed us to quantify animals' behavioural propensity to the conditioned corner.

We assumed that placing the animal in the cage context evokes the recall and possibly re-consolidation of the trained spatial-emotional association, consequently that this overall set-up will allow us to elucidate short- and long-term dynamics on a structural level.

\subsection{\label{sec:rg-simple} Remote recall involves smaller, more consolidated neuronal sub-populations in certain structures}

\begin{figure*}
 \centering
 \includegraphics[width=\textwidth]{cmb-fig-rr.pdf}
 \caption{\label{fig:rr} The differences in overall activity and activity co-localisation in investigated structures between rats with recent and remote recall.
 A. Comparison of the fraction of active nuclei (top) and co-localisation coefficient (bottom); both conditioned on the brain structure.
 In the remote recall, the activity has generally dropped in comparison to the recent recall, while the co-localisation has increased, which suggests the development of specialised neuron sub-populations.
 Box-plots adhere to a standard Tukey's definition, original data is superimposed as jittered points.
 B. Heatmap showing the values of brain activity descriptors found to significantly differ between groups in a machine learning-based multivariate analysis.
 Most straightforward effects were identified in the VTA and hippocampus, but also in cortical regions and amygdala.
 Values were shown as ranks for clarity, with rank one given to a lowest value (blue) and rank sixteen to the highest (red). 
 }
\end{figure*}
First, we investigated whether there are qualitative signs of strongly consolidated memory in the remote recall group, focusing on the most direct descriptors of inter-entry variability; the outcome of this analysis is summarised in Figure~\ref{fig:rr}A.

When comparing the overall brain activity in the remote group compared to the recent group, expressed as a percentage of any active nuclei, we see that there is a significant (p=0.04) drop of activity, conditioned over structures.
Moreover, this drop in overall activity was accompanied by an increase in co-localisation (p<0.001), suggesting an emergence of specialised neuronal sub-populations.
These differences were too subtle to be attributed to particular structures with standard statistical methods, however.

The activity was also fairly consistent between individual rats; the notable exception from the rule is nucleus accumbens, which noted striking individual variability in activity, which was not explained by the group. 
It can be explained by the animals’ behaviour, though.

\subsection{Recall of the remote memory has a distinct activity pattern from a recent recall}
The Boruta \cite{kursa2010feature} machine learning-based analysis of the differences between recent and remote recall has uncovered additional, more nuanced interactions, presented on Figure~\ref{fig:rr}B.

For VTA, it has detected the aforementioned pattern of higher overall activation in the recent group, yet more specialised in the remote group.
For hippocampus, the dentate gyrus area (DG) activity is higher in the recent recall group, and this effect is especially pronounced at the first entry.
On the other hand, in CA3 we see disjoint populations, active only in the first entry and indicating recent group, as well as active only in the second entry and indicating the remote group.
We have found that two cortical regions contain specific neuronal populations more active in the remote group; in the prelimbic cortex (PrL), it was active at the first entry, while in the agranular retrosplenial cortex (RSA), only at the second.

Two parameters of the amygdala activity at the first entry have been also found to be significant; overall activity of the lateral part and exclusive activity of the central part. 
Generally, they were higher in the recent group, but also in two rats from the remote group with slightly elevated hippocampal activity and high VTA co-localisation.


\subsection{High, memory remoteness-independent variation in NAcc is explained by behaviour}
\begin{figure*}
 \centering
 \includegraphics[width=\textwidth]{cmb-fig-cor.pdf}
 \caption{\label{fig:cor} The factors associated with the correctness in the first test entry (A) and second test entry (B).
 Heatmaps show the values of brain activity descriptors found to be significantly associated with correctness in a machine learning-based multivariate analysis.
 The high first-entry correctness score was related to nucleus accumbens and CA3 activity, while the central amygdala was a key driver in the second entry. 
 Values were shown as ranks for clarity, with rank one given to a lowest value (blue) and rank sixteen to the highest (red); gray encodes missing values. }
\end{figure*}

As mentioned in Section~\ref{sec:rg-simple}, there is a substantial variation of activity in certain structures which is not explained by the recent/remote group; in particular in the nucleus accumbens.
By using the trajectory data extracted from the video recordings of the behaviour, we have quantified the inclination of a rat to stay in the corner that was previously associated with the amphetamine injection.
To rule out the impact of variable exploration tendencies, we have expressed it as a correctness score, defined as a fraction of the total time spent in any corner which was spent in a conditioned one. 

While this score was not significantly correlated with the recent/remote recall group, we have identified significant molecular associations with machine learning; they are collected in Figure~\ref{fig:cor}.

High correctness was identified to be explained by the activity of hippocampal CA3 as well as both the shell and core of the nucleus accumbens.
Moreover, these activity levels were fairly consistent across two entries, henceforth almost all pattern parameters were selected.
A similar pattern applies to caudate putamen (CPu), yet to a lesser extent, because CPu activity at the second entry is less discriminating.

Finally, there are also significant correlations between first entry correctness and overall activity of the lateral amygdala (LA) as well as activity co-localisation in the granular retrosplenial cortex (RSG).

\subsection{Behaviour at each entry has distinct neuronal activity correlates}
While the overall activity in NAcc and CA3, which are strong correlates of correctness at the first entry, remains fairly consistent between entries, they cease to strongly explain the correctness at the second entry.
Only co-localisations at CA3 and NAcc core and overall activity of CA3 remain to be important.

On the other hand, the activity of the central amygdala (CeA) becomes a key driver, we also see significant interactions involving the infralimbic cortex (IL) and VTA. 
They are rather complex and involve a structure state at the first entry.

\subsection{Strongest inter-structural co-activation occurs at second entry}
\begin{figure}
 \centering
 \includegraphics[width=\columnwidth]{fig-cor-graph.pdf}
 \caption{\label{fig:gr-cor} Significant monotonic correlations between values of activity descriptors between investigated brain structures.
 Each panel presents graph for a certain activity descriptor; panels with no links (for an activity at any entry, \emph{any}, and at the second entry, \emph{2nd}) were omitted.
 The densest, non-trivial networks can be observed between sizes of neuron sub-populations active exclusively at certain entry.
 The weight of a link corresponds to an absolute value of Spearman correlation coefficient; black links represent positive, while red ones negative correlations. }
\end{figure}

We have investigated activity correlations across all investigated neuronal sub-populations in all structures.
The graphs constructed from these identified to be statistically significant ones are presented in Figure~\ref{fig:gr-cor} --- there were none such in the \emph{Any} and \emph{Second} classes.

This analysis has mostly uncovered trivial, inter-structural correlations, in particular within amygdala (BLA-LA-CeA), NAcc (shell \& core), retrosplenial cortex (RSG-RSA) and hippocampus (CA1-CA3, though not DG).

When analysing populations active exclusively during the first entry, we have found a significant negative correlation between CeA and PrL.
In comparison to the general first entry activity graphs, we see that both the internal coupling in NAcc and between CA1-CA3 have disappeared, which indicates that this synchronisation has persisted over both entries.
On the other hand, RSG-RSA link is present in both exclusive views but not in any more general one, suggesting it is a result of a more temporally localised phenomenon.

Most of the significant correlations identified are between the sizes of neuronal populations active exclusively during the second entry.
Here, we see intra-structural coherence in the amygdala, hippocampus, retrosplenial cortex and between prelimbic and infralimbic cortices, yet not in NAcc.
There are also inter-structural correlations, positive between the hippocampus, retrosplenial cortex and, to a lesser extent, amygdala; as well as negative correlations between the prelimbic cortex and both basolateral amygdala (BLA) and DG.

One should note, however, that the sample sizes are quite limited for a correlation network study, and this analysis is likely to have low sensitivity. 

\section{Discussion}
Contemporary models of memory generally involve multiple functional tiers.
Memories are first formed in short-term storage, which is transient as it must withstand a constant influx of new information.
To this end, encoded data is then selectively moved to long-term memory, based on a certain assessment of its possible usefulness --- this process may take substantial time, and it is generally faster when similar memories (often referred to as schemes) are already present in the long-term memory.

Long-term memory, on the other hand, is the first to react during recall, triggered by a stimulus consistent with a certain record.
The activation of short-term system follows, as recall almost always happens in circumstances where novel, relevant information can be acquired, and such would likely be used to update the recalled memory in a process of re-consolidation.

\subsection{Physiological changes consistent with memory consolidation}
It is reasonable to assume that certain memory utilises a progressively smaller and more consolidated population of neurons as it moves from a fresh, most verbose representation of current stimuli and emotional state to a consolidated, processed record.
This is indeed what we see when comparing rats with a remote recall from those with a recent recall; the observed activity is generally lower but more specialised.

Machine learning-based mining, capable of uncovering non-monotonic and multivariate interactions, has revealed a more detailed picture, showing significant differences on a particular structure level.
They were especially pronounced in the VTA and DG, but also in the amygdala, prelimbic and retrosplenial cortices, as well as in CA3.

The dynamics in the VTA were closest to the general trend of diminishing but specialising activity.
This structure is attributed to be an element of the reward system \cite{arias2010dopaminergic,polter2014stress} as well to be involved in memory processes \cite{cohen2022reward,diaz2021bidirectional}, especially when reward-related memories are considered.
Henceforth, the strong rewarding effect of amphetamine might have led to a formation of a specific neuronal population in this structure.

The higher activity of DG in the recent group is quite interesting, especially because it was also the least active part of the hippocampus; this may be connected with its speculated role in refining memories during encoding \cite{hainmueller2020dentate}. 
This process is expected to be more pronounced in the recent recall group, where animals experience a sudden break of the training routine.

Cortical activity patterns are particularly interesting, since they both consider an increase of activity in the remote recall group, suggesting that rats have in fact developed a specified neuron population during the hold off period.
They could not be detected by simple co-localisation analysis, however, because relevant cortical structures were not uniformly responding in either test entry --- in particular, said population in the prelimbic cortex was only active in the first entry, while this in RSA only in the second.

A similar argument can be made about CA3; the literature suggests it should be involved in the investigated phenomena, and indeed this is the case, yet its activity pattern goes beyond a simple consolidation model. 
Precisely, our results suggest that the recent recall group has a specific CA3 population active exclusively in the first entry, while the remote group has a similar population, yet active exclusively in the second entry.
A possible explanation of this pattern considers the fact that test sessions give an experience conflicting this from training --- amphetamine is not administered at all, not even in the conditioned corner.
This fact undoubtedly supports the dissolution of the trained association, which may be aided by CA3.
One can hypothesise that fresh memories are more volatile and a single conflicting observation is sufficient to trigger revision, while consolidated ones require stronger support and start to activate at the second entry.

The recent recall group was also characterised by an elevated activation of the lateral and central amygdala during the first entry; this is consistent with the aforementioned activity patterns of CA3 and VTA, leading to a conclusion that this trio was jointly involved in the encoding of freshly formed spatial-emotional association, utilising a network of projections well established in the literature. 
Our results suggest that this action is transient and can be easily diminished by conflicting experiences, though.

Interestingly, clustering based on ML-selected parameters has uncovered two clear outliers (a05 and a02) which, despite being in a recent recall group, exhibited activity patterns of the remote group.
We interpret this as a sign of early consolidation, possibly triggered by an earlier spatial memory or individual differences in emotional state or a reaction to amphetamine.

\subsection{Utilisation of a memory depends on different factors than its consolidation}
We have used a correctness score to quantity the animal's propensity to the corner associated with the amphetamine injection, which we believe corresponds to goal-directed drug seeking.
There is no group effect in this parameter, which we interpret as an indication that all animals have correctly recalled the spatial organisation of the train/test cage and identified the conditioned corner, yet adjusted the seeking and anticipatory behaviours based on other factors.
In particular, this involves both areas of NAcc, which express higher activity in rats with a larger initial correctness score; this is also consistent with a very high individual variability of NAcc activity patterns.

NAcc is a key hub in the cortico-limbic circuitry directing action selection and decision-making in reward-related tasks \cite{floresco2018differential}, and its activity is bound to the expression of emotion, in particular, reflected by ultrasonic vocalisation 
\cite{hamed2016non,sohn2022linking,brudzynski2012quinpirole,burgdorf2001nucleus}, and with processing current values and reward prediction \cite{humphries2010ventral,bornstein2011multiplicity,khamassi2012integrating,martin2000effects}.
To this end, we conclude that the experiment likely captured activation of NAcc neurons positively associated with reward anticipation, which was a reaction to the recognition of substance administration context, possibly induced directly by cortex through well-established connections \cite{groenewegen1982cortical,jayaraman1985organization,powell1976connections,zahm1992significance}.

The strength of this action was shaped by the individual variability among rats, however, which caused different behavioural outcomes.
The very origin of this variability is unclear, yet it likely arises at a network level because of the aforementioned hub nature of NAcc.
In particular, the first-entry correctness is also positively correlated with the activity of the caudate putamen and hippocampus CA3, as well as, to a lower extent, with the overall activity of amygdala LA and activity co-localisation in RSG.

It is crucial to note that the NAcc activity persisted between test entries, despite the fact that it was not indicative of the correctness in the second entry.
In general, we have not assumed both test sessions to be equal, as the first one establishes a novel and conflicting context for the conditioned corner, as there is no amphetamine injection consistent with previous experience.
This is directly visible on Figure~\ref{fig:disc}, which summarises all the machine learning results.

While the overall NAcc activity has not diminished due to this fact yet the behaviour has changed, we believe this is a consequence of a tertiary mechanism that has modulated the NAcc influence.
The machine learning analysis of the correlates of the second entry correctness scores suggests that this can be attributed to an aforementioned CA3-amygdala-VTA trio.
In particular, a higher score is predicted by activation of the CeA, but during the first entry; this can be attributed to a hypothesis that the sustenance of the anticipatory actions is promoted by the original emotional arousal. 
CA3 and VTA activity effects are more complex and require further insights.

\section{Conclusions}
\begin{figure*}
 \centering
 \includegraphics[width=\textwidth]{fig-sg.pdf}
 \caption{\label{fig:disc} Overview of the overlap between machine learning-selected significantly important factors connected with the memory recall group and correctness score in either test entry. 
 Although hippocampus, amygdala and nucleus accumbens re-appear in all contexts, their involvement differs at the level of finer-grained neuronal populations.
 }
\end{figure*}
Inspired by theoretical models of memory dynamics, we formed a hypothesis that memory consolidation involves the emergence of specialised neuronal populations.
To this end, we have scanned the brain for traces of such, i.e., regions which exhibit lower but more co-localised activity when comparing rats with maturated memories to freshly trained ones, within structures connected with memory functions and emotional processing. 
We have confirmed that pattern on the whole-brain level, as well as that it is most pronounced in the VTA.

The dynamics of cortical regions, generally regarded to be the prime destination for consolidating memories, proved to be more complex.
This may be a sign of higher-order phenomena triggered by an asymmetry between training and testing conditions, namely a need to re-encode the spatial memory with a novel, conflicting experience acquired in testing.
This process appears to involve hippocampus CA3 and causes increasing intra-structural coordination, but further insight is required to verify these observations.

Finally, we have shown that the behavioural outcome induced by memory recall can be heavily modulated by independent factors, including individual variability, especially when emotional processing is substantially involved --- in particular, we have found nucleus accumbens to be a key structure enforcing action, while the amygdala to  likely be either an integrator or a modulator.
Anyhow, molecular and physiological observations provide a more direct and reliable quantification of the properties of memories.
On the other hand, considering individual variability is crucial for the development of effective interventions, both experimental and therapeutic.

\section*{Funding}
R.C., M.F. and A.H. were supported by grant Sonata-bis 2014/14/E/NZ4/00172 from the National Science Centre Poland.
A.H. and W.K. were supported by grant Opus 2018/29/B/NZ7/02021 from the National Science Centre Poland.
 
\section*{Acknowledgments}
We wish to thank Tomasz Gomoła for the construction a 3D model of an experimental setup.

\section{Methods}

\subsection{Animals}

Adult male Long-Evans rats ($n=18$, $180\pm20g$) were used in the experiment.
Animals were purchased from a licensed breeder (the Polish Academy of Science Medical Research Center, Warsaw, Poland), and housed in standard laboratory conditions under $12h$:$12h$ light:dark cycles (lights on at 7 a.m.), at a constant temperature ($21\pm 2 \degree C$) with $70\%$ humidity. 
Rats had free access to both food and water. 
All experiments were performed in accordance with the European Communities Council Directive of 24 November 1986 (86/609 EEC).
Local Ethical Committee approved all experimental procedures using animal subjects (539/2018). 


\subsection{Behavioural experiment}
All experiments utilised the same $1m \times 1m$ cage, partitioned into 5 regions (four corners and a central interconnecting space) with translucent walls equipped with doors.
In the \emph{training} configuration doors are closed, henceforth a rat can be confined in a selected corner but retains the capacity to observe the whole cage through the wall.
On the other hand, in the \emph{testing} configuration, doors are opened, and a rat can freely roam around the cage.
The whole setup is softly illuminated by an overhung asymmetric array of static colour lights, which serves as a visual-spatial cue.
Otherwise, corners were identical in terms of surface texture, colour or size; the cage was cleaned with $70/100$ ethanol solution before each rat placement to remove dirt and scent marks. 

Animals were first habituated to the experimenter and cage; in this phase, doors were opened and the visual cue array was inactive.
After habituation, the cage was switched to the training configuration, and animals were trained in the following manner.
In each of the six training days, the animal was placed confined in a corner c1, c3, c2, c3, c4 and c3, respectively, for 15 minutes. 
During this time, the animal was injected with either amphetamine at a dose of 1.5$mg/kg$, when in corner c3, or saline (1$ml/kg$; i.p.), in other corners; this way, c3 was the conditioned corner.

After training, animals were, according to the pre-assigned group, either immediately transferred to testing (recent group), or first underwent a hold-off in their home cage for 14 days (remote group), and then tested.
The testing procedure was identical for all animals and was performed as follows.
The animal was placed in the centre of the cage in the testing configuration, with doors opened.
Then, it was allowed to freely move and explore for five minutes; in particular, the rat could visit the conditioned corner.
After this first entry, animals were put on hold in a separate cage for 20 minutes.
Finally, animals were placed again in the centre of the cage for another 5 minutes, which constituted the second entry.

Immediately after testing, each animal was decapitated under isoflurane anaesthesia; its brain was isolated and flash-frozen in a dry-ice cooled isopentane bath.
Afterwards, brains were awaiting catFISH reaction stored aluminium foil-wrapped in a deep freezer set to -80$\degree C$.

All sessions (habituation, six training entries and two test entries) took place under video surveillance; recordings of the test sessions were later used to reconstruct rat trajectories using the DeepLabCut toolbox \cite{mathis2018deeplabcut}.

\subsection {Fluorescent staining of IEGs}
To optimise the slicing process, we arranged brains side by side in blocks of four, embedded in a medium with an optimal slicing temperature (OCT; Sakura).
The blocks were cut in a cryostat (Leica CM 1850, Germany) into 20$\mu m$ sections, which were mounted on gelatin-coated SuperFrost slides (ThermoFisher).

Next, slices undergo fluorescent staining according to an established protocol \cite{vazdarjanova2004differences,chawla20043d, kubik2012loss}.
Fluorescein- and digoxigenin-labelled antisense riboprobes for 3’UTR of, respectively, Homer 1a and Arc mRNA were applied to slices and allowed to complete hybridisation overnight, in a single step.
Then, both kinds of probes were sequentially detected, first with anti-fluorescein and then with anti-digoxigenin.
Homer 1a probes were stained with a tyramide-fluorescein signal amplification system (TSA-Fluorescein) and Arc probes with TSA-Cy3 (Perkin-Elmer).
Furthermore, the slides were also incubated with DAPI nuclear counter-stain (Invitrogen, ThermoFisher), to visualise the nuclei.
Finally, prepared slides were cover-slipped with an anti-fade media (Vectashield, Vector labs) and sealed with nail polish.


\subsection{Image acquisition and analysis}
Stained slices containing representative samples of interesting structures were scanned using Inverted Leica DMI 6000 microscope with Andor DSD2 Confocal Module and HC PL APO CS2 20x/0.70 Immersive objective. 
Fluorescence was excited with 200W halogen lamp.
To detect nuclei, Homer-1a and Arc, three fluorescent filtersets were used, respectively: DAPI (exc. 390$\pm$40$nm$, 405 DM, em. 452$\pm$45$nm$), FITC (exc. 483$\pm$28$nm$, 488 DM, em. 525$\pm$45$nm$) and Cy3 (exc. 556$\pm$20$nm$, 561 DM, em. 609$\pm$54$nm$).


We optimised the settings to obtain bright, intranuclear foci of ongoing IEG transcription.
The laser power, gain, and offset, as well as exposure times, were always set for the whole slide to ensure the best possible signal without substantial spatial variation.  
The scan was repeated to construct a z-stack; we rejected three bottom- and top-most layers so that only whole cells were considered later.
Signals from individual filters were routed into separate channels of the final image.

For each structure, we performed computer-aided manual alignment of a brain atlas to the sub-sampled, flattened images of appropriate slices (Figure~\ref{fig:scheme}E); this way, we have defined numerically defined ROIs for further automatic analysis.

Next, for each ROI, we used a computer vision pipeline to interpret the geometry of the distribution of reporter proteins.
We have pooled each structure from both cerebral hemispheres.
In particular, for FITC and DIG/Cy3 emission channels, we have used the ICY dot-finder to identify localised aggregations of the reporter, marking the locations of the mRNA particle of the corresponding IEG.
For the nuclei identification, we have fitted ellipses to a coherent circular bright patch in the DAPI emission channel, using a custom code based on the watershed algorithm \cite{ng2006medical}.
Finally, for each nucleus region, we counted the number of dots for either IEG; this way, each ROI got quantified as fractions of nuclei of four classes: both Arc and Homer-1a positive, Arc-only positive, Homer-1a only positive, and negative.
This analysis was repeated for each z-layer, and we collected the median values of said fractions as a final result.

Additionally, we have calculated the co-localisation coefficient, given as a fraction of both Arc and Homer-1a positive nuclei divided by its expected value under the independent placement hypothesis, which is a product of fractions of Arc and Homer-1a positive nuclei. 
This coefficient is positive but unbounded and is expected to be one given the independence of neuron activation in both entries, under one when the activation is somehow exclusive to either entry and finally over one when neuron activations in both entries are correlated.

\subsection{Statistical analysis}
For a basic comparison of the activity between the recall groups, we have used a stratified, two-sided Mann-Whitney-Wilcoxon test.
The Spearman correlation test was used for the reconstruction of inter-structural interaction graphs.
We have applied 0.05 as a significance threshold for p-values, and use Holm's correction \cite{holm1979simple} for multiple testing, expect in the cross-structure correlation analysis which used Benjami-Hochberg FDR correction.

The Boruta method \cite{kursa2010feature} was applied for the machine learning search for multivariate and non-linear interactions, using the standard Random Forest importance source with 50 000 trees wrapped into the impute transdapter; tentative selections were collected together with the confirmed ones.
To stabilise the results, we repeated this procedure thirty times and finally reported features selected in at least half of them.

\section*{Data availability}
Raw data are available on Mendeley Data:\\ \verb|http://dx.doi.org/10.17632/3fg8khy6dp.1|

\section*{Code availability}
Code for reproducing data analysis is available on GitLab: \\ \verb|https://gitlab.com/neuro-reward/reward-memory|

\bibliography{catfish}


\end{document}
