## Supplementary figures and images for "Spatio-temporal mechanisms of consolidation, recall and reconsolidation in reward-related memory trace"

### cmb-fig-cor.pdf

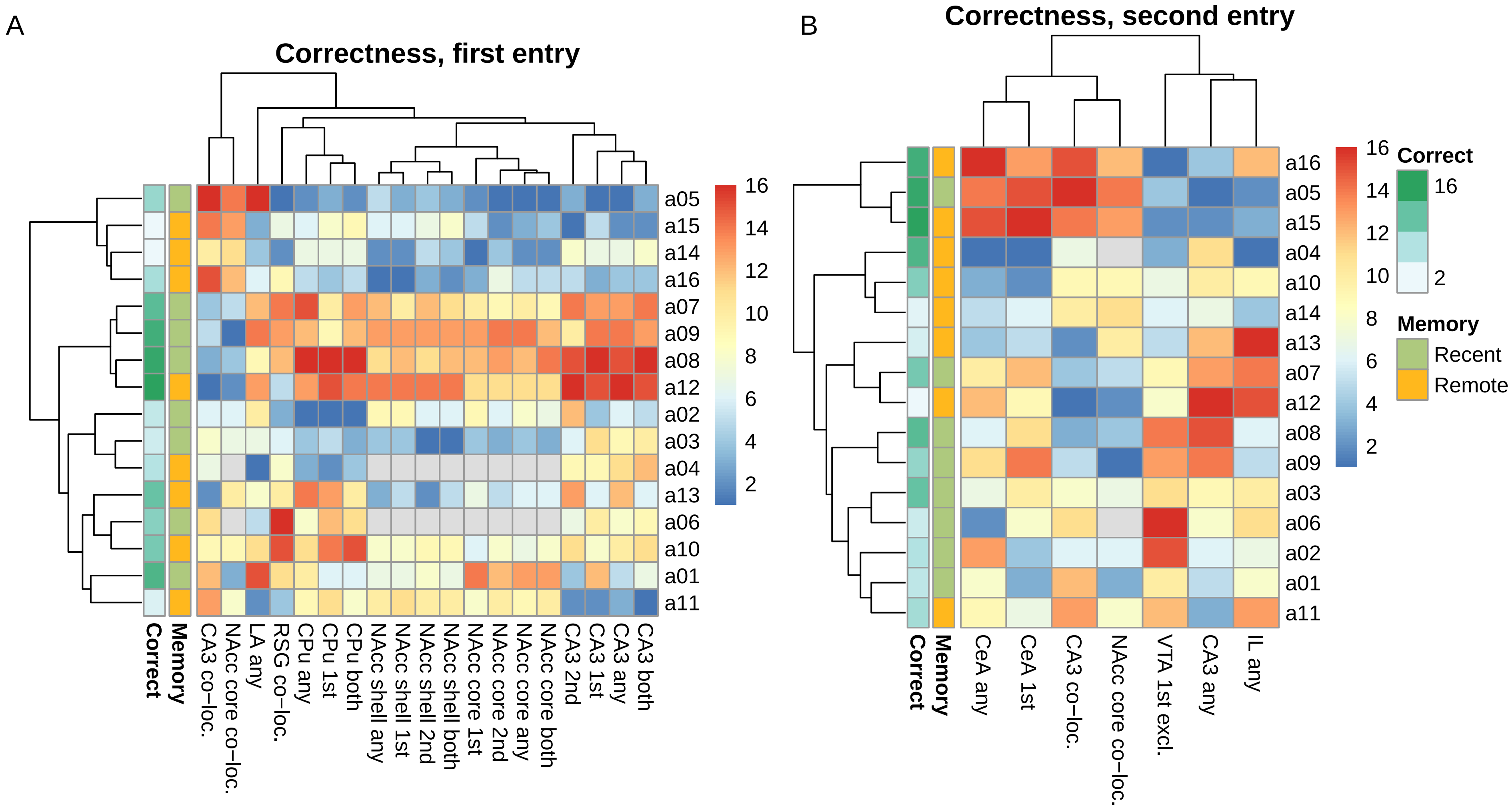

### cmb-fig-rr.pdf

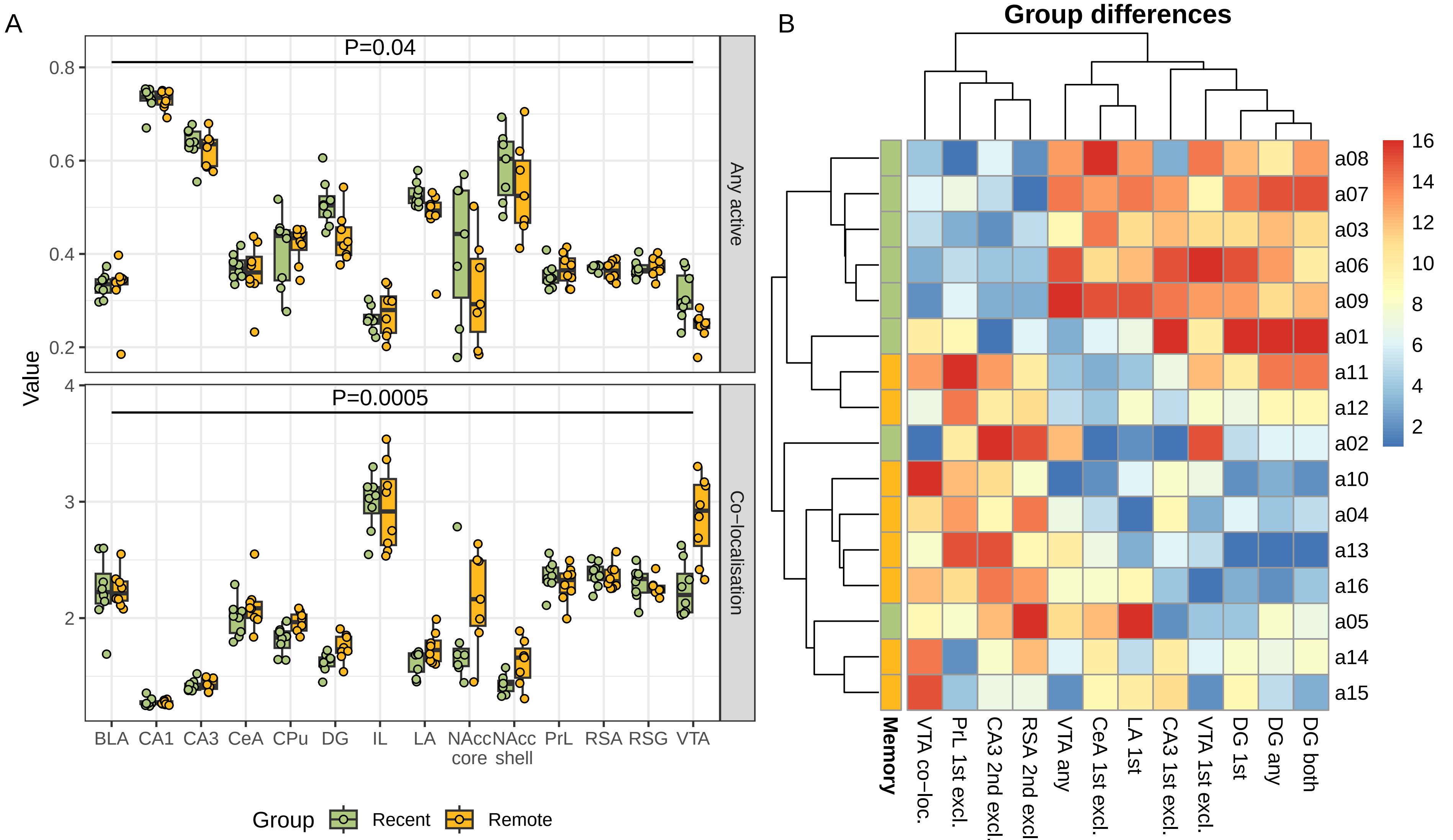

### fig-cor-graph.pdf

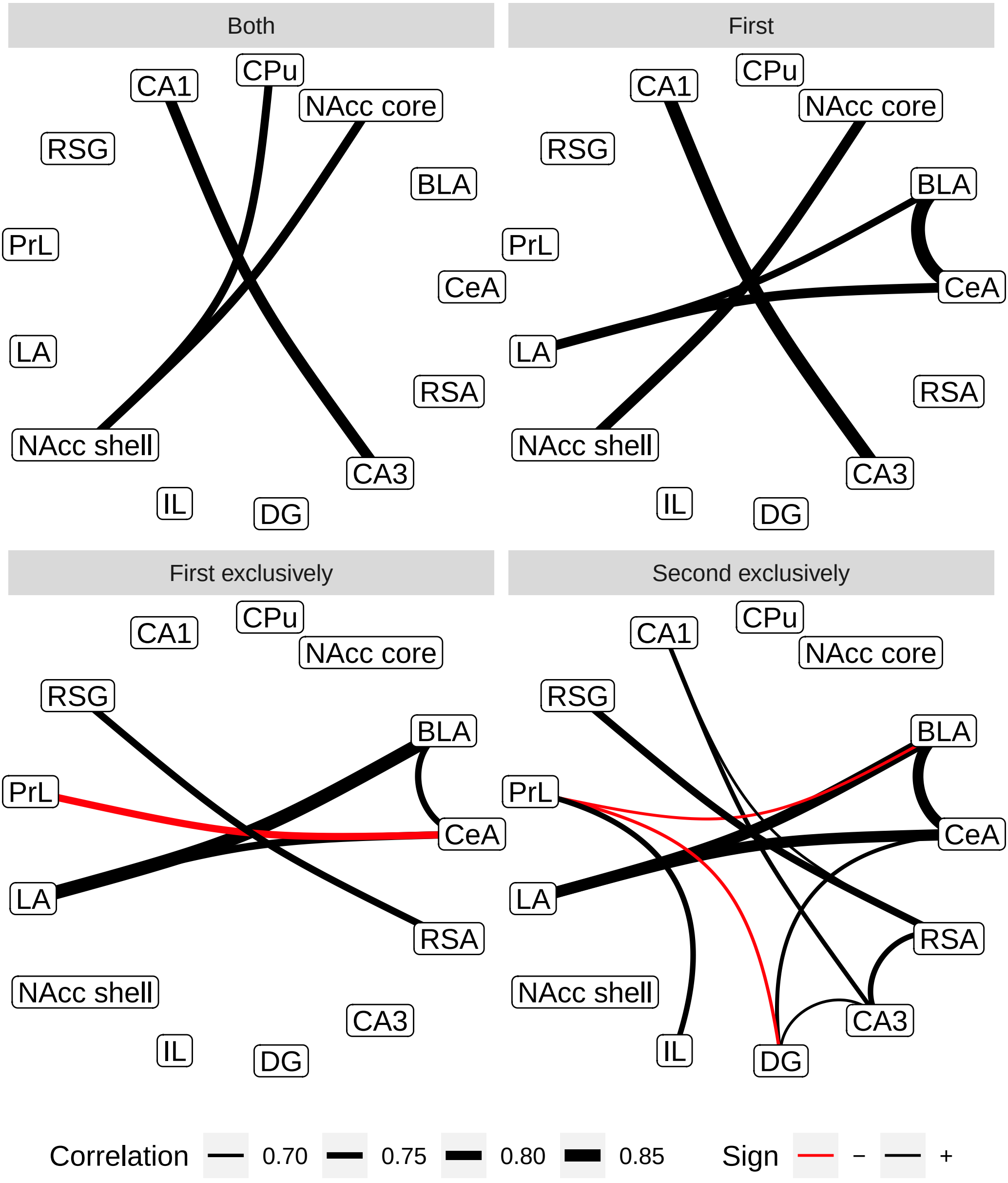

### fig-sg.pdf

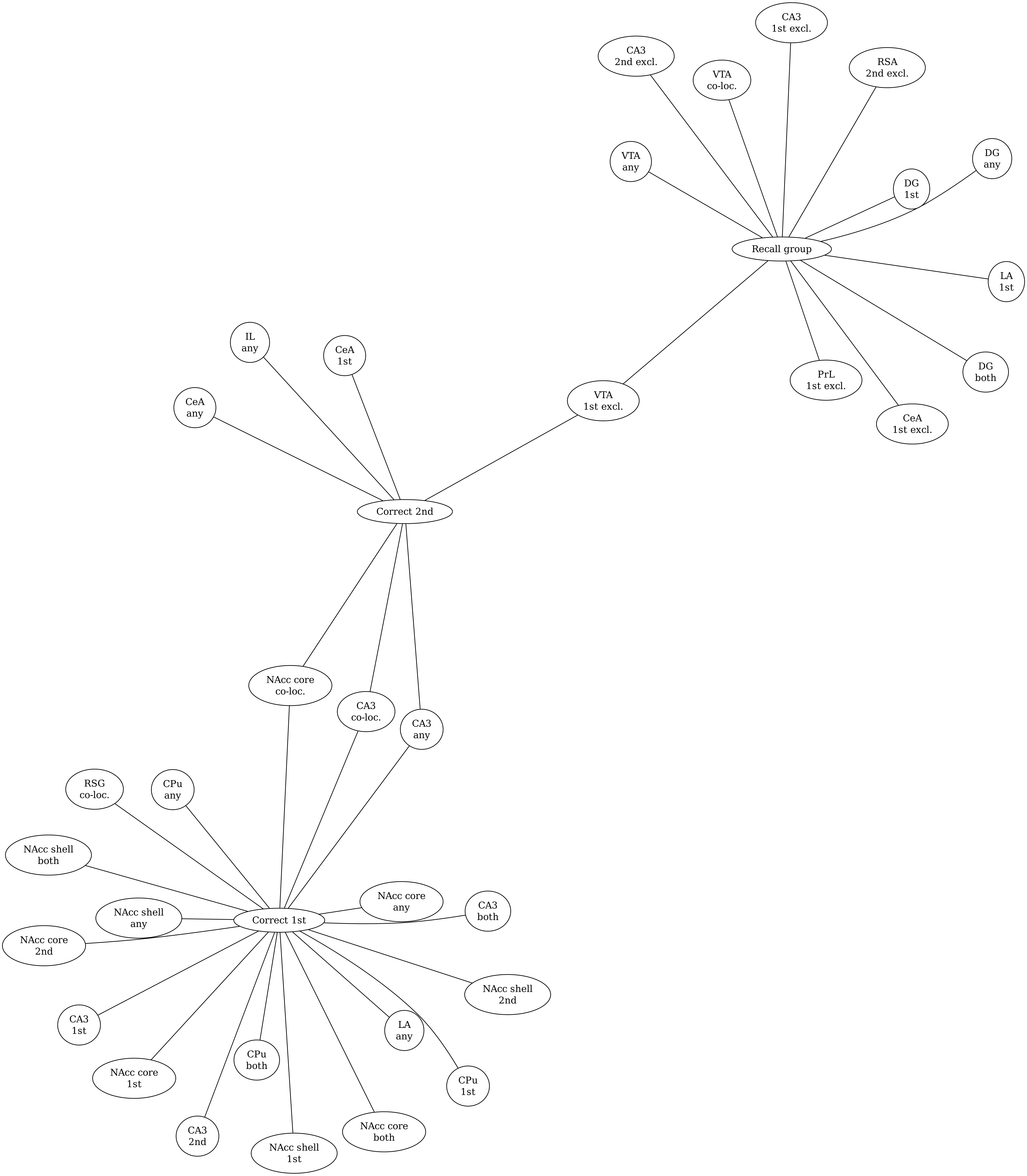

### scheme.pdf

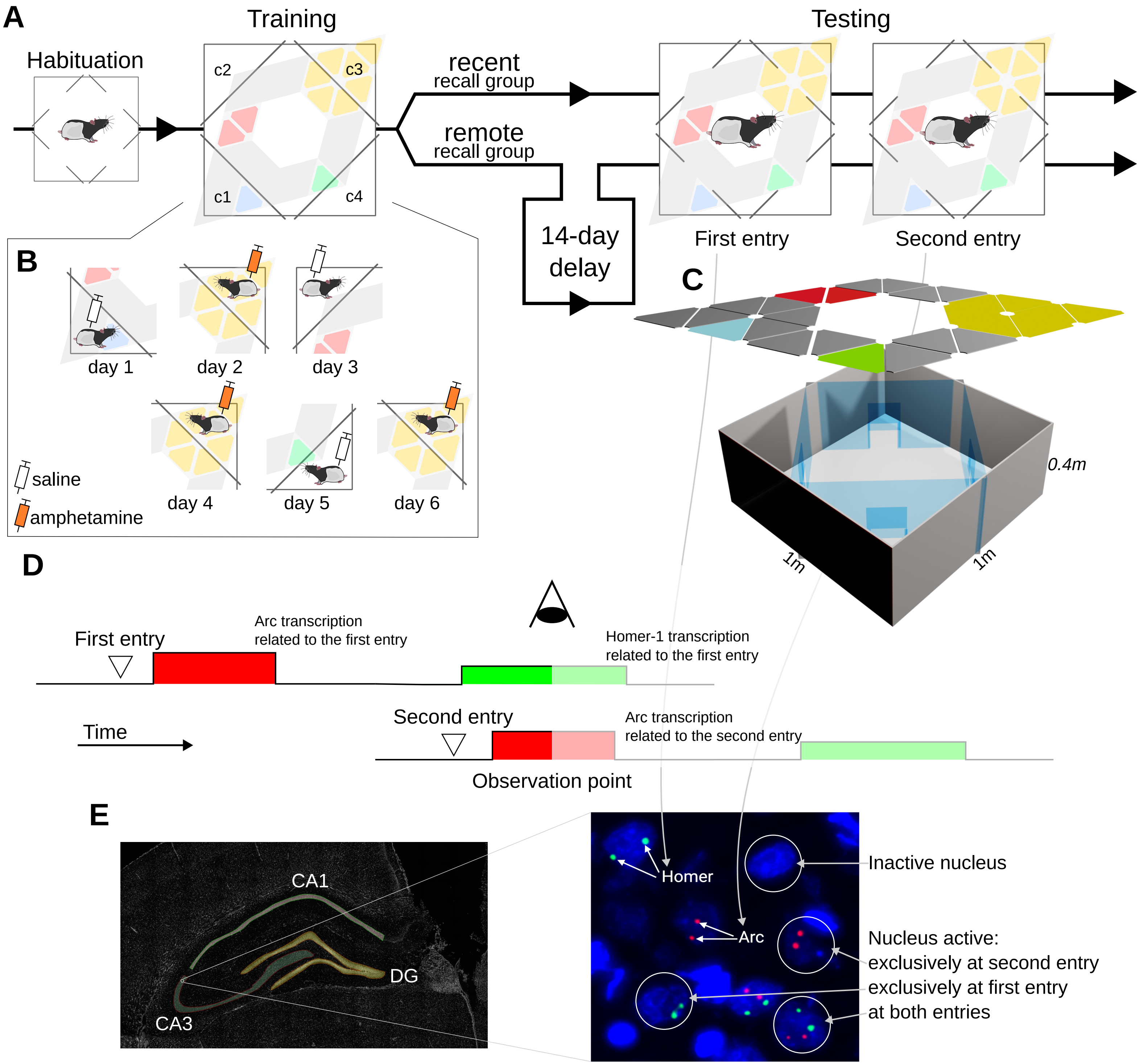
